# Interleukin-6 mediates PSAT1 expression and serine metabolism in TSC2-deficient cells

**DOI:** 10.1101/2021.05.17.444471

**Authors:** Ji Wang, Harilaos Filippakis, Thomas Hougard, Heng Du, Chenyang Ye, Heng-Jia Liu, Long Zhang, Khadijah Hindi, Shefali Bagwe, Julie Nijmeh, John M. Asara, Wei Shi, Souheil El-Chemaly, Elizabeth P. Henske, Hilaire C. Lam

**Affiliations:** Pulmonary and Critical Care Medicine, Brigham and Women’s Hospital, Harvard Medical School, Boston, MA 02115; The F.M. Kirby Neurobiology Center, Boston Children’s Hospital, Department of Neurology, Harvard Medical School, Boston, MA 02115; Division of Signal Transduction, Beth Israel Deaconess Medical Center and Department of Medicine, Harvard Medical School Boston, MA, USA 02215; Department of Surgery, Children’s Hospital Los Angeles, Keck School of Medicine, University of Southern California, Los Angeles, CA 90027

**Keywords:** Tuberous sclerosis complex (TSC) / Lymphangioleiomyomatosis (LAM), mTORC1/ Rapamycin, Interleukin-6 (IL-6), serine metabolism, phosphoserine aminotransferase 1 (PSAT1)

## Abstract

Tuberous sclerosis complex (TSC) and lymphangioleiomyomatosis (LAM) are caused by aberrant mechanistic Target of Rapamycin Complex 1 (mTORC1) activation due to loss of either *TSC1* or *TSC2*. Cytokine profiling of TSC2-deficient LAM patient-derived cells revealed striking upregulation of Interleukin-6 (IL-6). LAM patient plasma contained increased circulating IL-6 compared with healthy controls, and TSC2-deficient cells showed upregulation of IL-6 transcription and secretion compared to wildtype cells. IL-6 blockade repressed the proliferation and migration of TSC2-deficient cells and reduced oxygen consumption and extracellular acidification. U-^13^C glucose tracing revealed that IL-6 knockout reduced 3-phosphoserine and serine production in TSC2-deficient cells, implicating IL-6 in *de novo* serine metabolism. IL-6 knockout reduced expression of phosphoserine aminotransferase 1 (PSAT1), an essential enzyme in serine biosynthesis. Importantly, recombinant IL-6 treatment rescued PSAT1 expression in the TSC2-deficient, IL-6 knockout clones selectively and had no effect on wildtype cells. Treatment with anti-IL-6 (aIL-6) antibody similarly reduced cell proliferation and migration and reduced renal tumors in *Tsc2*^*+/-*^ mice, while reducing PSAT1 expression. These data reveal a novel mechanism through which IL-6 regulates serine biosynthesis, with potential relevance to the therapy of tumors with mTORC1 hyperactivity.

**Classification:** Major category: Biological Sciences Minor category: Cell Biology

## Introduction

Tuberous Sclerosis Complex (TSC) is an autosomal dominant tumor suppressor syndrome that affects one in 10,000 infants (1-4). The majority of patients suffer from neurodevelopmental conditions including epilepsy, autism and cognitive impairment. Neoplastic lesions in the brain, skin, heart, kidney and lung are the primary causes of patient morbidity and mortality, particularly later in life (5, 6). Renal angiomyolipomas affect ∼70% of patients by 10 years of age (7). Lymphangioleiomyomatosis (LAM), characterized by pulmonary nodules and irreversible progressive cystic lung destruction, almost exclusively affects females with TSC and can also affect women with sporadic LAM (4, 8).

TSC is caused by inactivating mutations in *TSC1* or *TSC2*, resulting in aberrant activation of mechanistic Target of Rapamycin Complex 1 (mTORC1), a master regulator of cellular metabolism (9). Constitutive mTORC1 activation leads to extensive changes in signaling pathways and promotes lipid, nucleotide and protein biosynthesis contributing to the dysregulated growth and proliferation of cells in patients with TSC (10, 11). mTORC1 can be directly targeted by the allosteric inhibitor rapamycin and related analogs, “rapalogs,” as well as catalytic mTOR inhibitors. Since mTORC1 inhibition primarily exerts cytostatic effects (7, 12), identifying novel therapeutic targets that can yield more durable or cytotoxic clinical responses is a key focus of ongoing TSC research.

Interleukin-6 (IL-6) is a secreted cytokine and critical mediator of inflammation (13). IL-6 binds to either soluble or membrane bound IL-6 receptor α. The IL-6/IL-6Rα complex then interacts with gp130 to activate signaling via Janus Kinase / Signal Transducer and Activator of Transcription 3 (JAK/STAT3). STAT3 regulates the transcription of hundreds of genes, including IL-6. This positive feedback loop has been previously shown to be an epigenetic mechanism of transformation downstream of transient Ras activation (14). STAT3 activation by mTORC1 is a well described feature of TSC lesions, along with an increase in IL-6 production (15-23).

In support of prior findings, we find that IL-6 is upregulated in the plasma of patients with LAM and in preclinical models of TSC, in a TSC2- and mTORC1-dependent manner. By inhibiting IL-6 both genetically and with neutralizing antibodies, we find that TSC2-deficient cells depend on IL-6 to support the cell-autonomous metabolic reprogramming necessary for proliferation and migration. In particular, we implicate IL-6 in the regulation of *de novo* serine synthesis in TSC2-deficient cells. The enzymatic reactions necessary for *de novo* serine metabolism produce serine, as well as antioxidants and the tricarboxylic acid (TCA) cycle intermediate, a-ketoglutarate. Specifically, the first rate-limiting step is executed by phosphoglycerate dehydrogenase (PHGDH), which converts the glycolytic intermediate 3-phoshphogylcerate to 3-phosphohydroxypyruvate, regenerating NADH from NAD. The next step is mediated by phosphoserine aminotransferase 1 (PSAT1), which converts 3-phosphoydroxypyruvate to 3-phosphoserine by transferring the amino group from glutamate and producing the TCA cycle intermediate a-ketoglutarate. Finally, 3-phosphoserine is converted to serine by phosphoserine phosphatase (PSPH). Serine can also be generated from glycine via the reversible action of serine hydroxymethyltransferase (SHMT) enzymes. Recent reviews have highlighted the importance of *de novo* serine metabolism in cancer survival and progression (24, 25). Our data implicate IL-6 as a novel regulator of *de novo* serine metabolism in mTORC1 hyperactive cells, thereby promoting the tumorigenic potential of TSC2-deficient cells.

## Results

### IL-6 is upregulated in LAM patient plasma and preclinical LAM and TSC models

Previous studies have shown that TSC2-deficient cells have a unique secretome, which may support the proliferation and metastatic potential of TSC tumors and LAM nodules by both cell autonomous and paracrine effects (21-23, 26). We performed a cytokine array using the TSC2-deficient 621-101 cell line, derived from a human angiomyolipoma, in comparison to the human embryonic kidney cell line, HEK293. The most robustly upregulated cytokine in 621-101 cells was IL-6 (**Fig. S1A-D**), a factor previously reported in LAM *in vivo* models and patient-derived cells (21, 23). We discovered that IL-6 is also upregulated in plasma from LAM patients compared to healthy controls (**Fig. 1A**).

**Figure 1.**
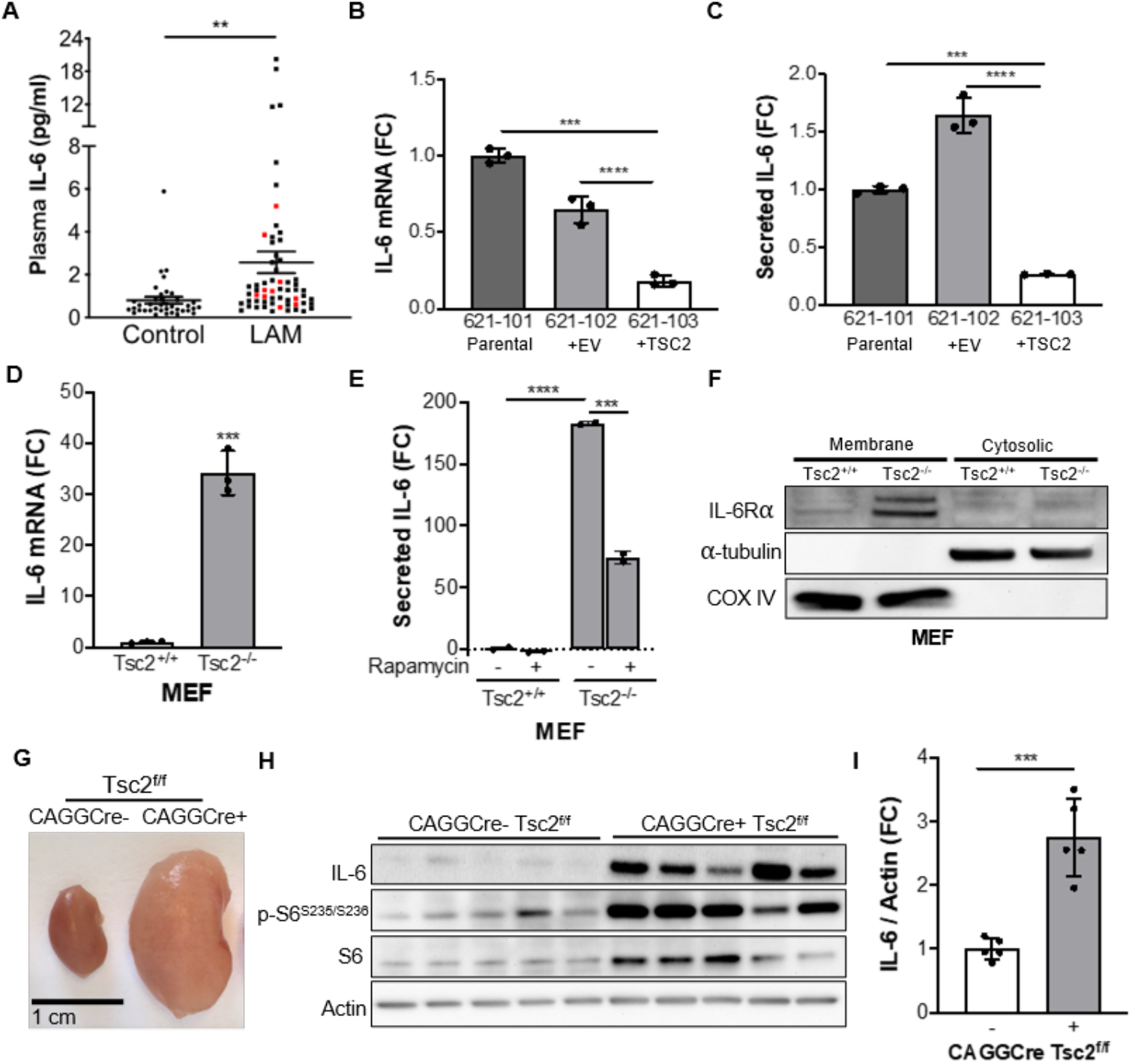
IL-6 is overexpressed in TSC2-deficient cells and tissues. (*A*) IL-6 was increased in the plasma of LAM patients (total n=60, black dots: sporadic-LAM, n=50; red dots: TSC-LAM, n=10) compared to healthy control (n=38). The data are presented as the mean ± SEM. (*B*) IL-6 mRNA expression by qRT-PCR and (*C*) secreted IL-6 measured by ELISA are increased in TSC2-deficient human angiomyolipoma parental cells (621-101) and cells expressing empty vector (621-102), compared to cells with TSC2 addback (621-103). (*D*) qRT-PCR of IL-6 mRNA expression showing a 30-fold increase in Tsc2^-/-^ MEFs compared to Tsc2^+/+^ MEFs. (*E*) Rapamycin treatment decreases secreted IL-6 in TSC2-deficient MEFs as measured by ELISA, (rapamycin; 20nM, 24 hours). (*F*) Western blot of membrane and cytosolic protein fractions of TSC2-deficient and wildtype MEFs. IL-6 receptor (IL-6Rα) is highly expressed in TSC2-deficient MEFs compared to TSC2-expressing MEFs. Cells were cultured in serum-free DMEM for 24 hours before harvesting. α-tubulin used as a cytosolic fraction marker and COX IV as a membrane fraction marker. (*G*) Representative kidneys of *CAGGCre-ER*^*TM+/-*^; *Tsc2*^*f/f*^ and *CAGGCre-ER*^*TM-/-*^; *Tsc2*^*f/f*^ mice. (*H*) Western blot of *CAGGCre-ER*^*TM+/-*^; *Tsc2*^*f/f*^ and *CAGGCre-ER*^*TM-/-*^; *Tsc2*^*f/f*^ kidneys showing increased IL-6 expression upon TSC2 loss. (*I*) Densitometry of IL-6 protein levels normalized to actin in *CAGGCre-ER*^*TM+/-*^; *Tsc2*^*f/f*^ and *CAGGCre-ER*^*TM-/-*^; *Tsc2*^*f/f*^ kidney lysates shown in *H*. Data are presented as the mean ± SD of three independent experiments, unless indicated otherwise. One-Way ANOVA, Two-Way ANOVA, or Student’s t test were used for statistical analysis. *p < 0.05, **p < 0.01, ***p < 0.001, ****p < 0.0001.

We next confirmed that IL-6 expression is TSC2-dependent by comparing empty vector (621-102) or TSC2-addback cells (621-103) derived from the parental angiomyolipoma 621-101 cell line. TSC2 re-expression significantly reduced *IL-6* mRNA expression and secretion of IL-6 (∼70%, p<0.0001) compared to the TSC2-deficient lines (**Fig. 1B and *C***). We next determined the expression and secretion of IL-6 in two additional pairs of TSC2-deficient and expressing cell lines: TTJ cells, derived from a renal tumor of a *Tsc2*^*+/-*^ mouse, expressing either empty vector or re-expressing TSC2, and *Tsc2*^*+/+*^ and *Tsc2*^*-/-*^ mouse embryonic fibroblasts (MEFs) (27, 28). IL-6 mRNA expression was upregulated by 7-fold in the TSC2-deficient TTJ cells relative to TSC2-expressing control cells (p<0.01, **Fig. S1E**) and by ∼30-fold in the TSC2-deficient MEFs relative to TSC2-expressign MEFs (p<0.001, **Fig. 1D**). Secretion of IL-6 was also increased in the TTJ cells (p<0.05, **Fig. S1F**) and TSC2-deficient MEFs (p<0.0001, **Fig. 1E**). Rapamycin (20nM, 24h) partially reduced IL-6 secretion by ∼50% (p<0.001) in TSC2-deficient MEFs (**Fig. 1E**).

Importantly, we found that IL-6 receptor α (IL-6Rα) is upregulated in the membrane fraction of TSC2-deficient compared to TSC2-expressing MEFs, suggesting that IL-6 can act in an autocrine manner (**Fig. 1F**). IL-6 expression was also significantly elevated ∼2.5-fold (p<0.001) in kidney homogenates of *CAGGCre-ER*^*TM+/-*^; *Tsc2*^*f/f*^ mice, which are characterized by cystic kidney disease driven mTORC1 hyperactivation, compared to *CAGGCre-ER*^*TM-/-*^; *Tsc2*^*f/f*^ control mice (**Fig.1*G*-*I***) (29, 30).

In summary, these data show that IL-6 is upregulated in patient plasma and consistently across numerous *in vitro* and *in vivo* models of TSC and LAM. Furthermore, IL-6 expression is both TSC2- and mTORC1-dependent. Finally, the increased expression of IL-6Ra on the TSC2-deficient cells suggests that IL-6 may be secreted and detected by TSC2-deficient cells thereby exerting cell autonomous effects.

### IL-6 knockout suppresses proliferation, migration, and induces a metabolic quiescent state in TSC2-deficient cells

To investigate the dependance of TSC2-deficient cells on IL-6, we used CRISPR/Cas9 to knockout IL-6 from TSC2-deficient MEFs. We validated the knockout by measuring IL-6 secretion in three separate single cell clones generated from a single CRISPR/Cas9 guide (**Fig. 2*A***). Genetic knockout of IL-6 decreased the proliferation of TSC2-deficient cells in serum-free conditions in comparison to IL-6 expressing TSC2-deficient cells (>30%, p<0.001) as assessed by crystal violet staining as an indicator of cell density changes over 3 days (**Fig. 2*B***). In order to maximize the differences between TSC2-deficient and wildtype control cells and to eliminate the possibility of the cells reacting to bovine IL-6 in the serum, we chose serum-free conditions for all subsequent experiments. Since metastasis is a key aspect of LAM pathogenesis, we also wanted to investigate the impact of IL-6 on the migration capacity of TSC2-deficient cells (31, 32). We discovered that IL-6 knockout also suppressed migration of TSC2-deficient cells through transwells towards a chemoattractant and in wound healing assays compared to control cells (**Fig. 2*C, D*, and S2*A, B***). Importantly, conditioned media from TSC2-deficient cells with intact IL-6 rescued proliferation of the IL-6 knockout cells (**Fig. S2*C***). Treatment with rIL-6 (200 pg/ml) rescued the proliferation and migration of the TSC2 deficient cells with IL-6 knockout (**Fig. S2D and E**)

**Figure 2.**
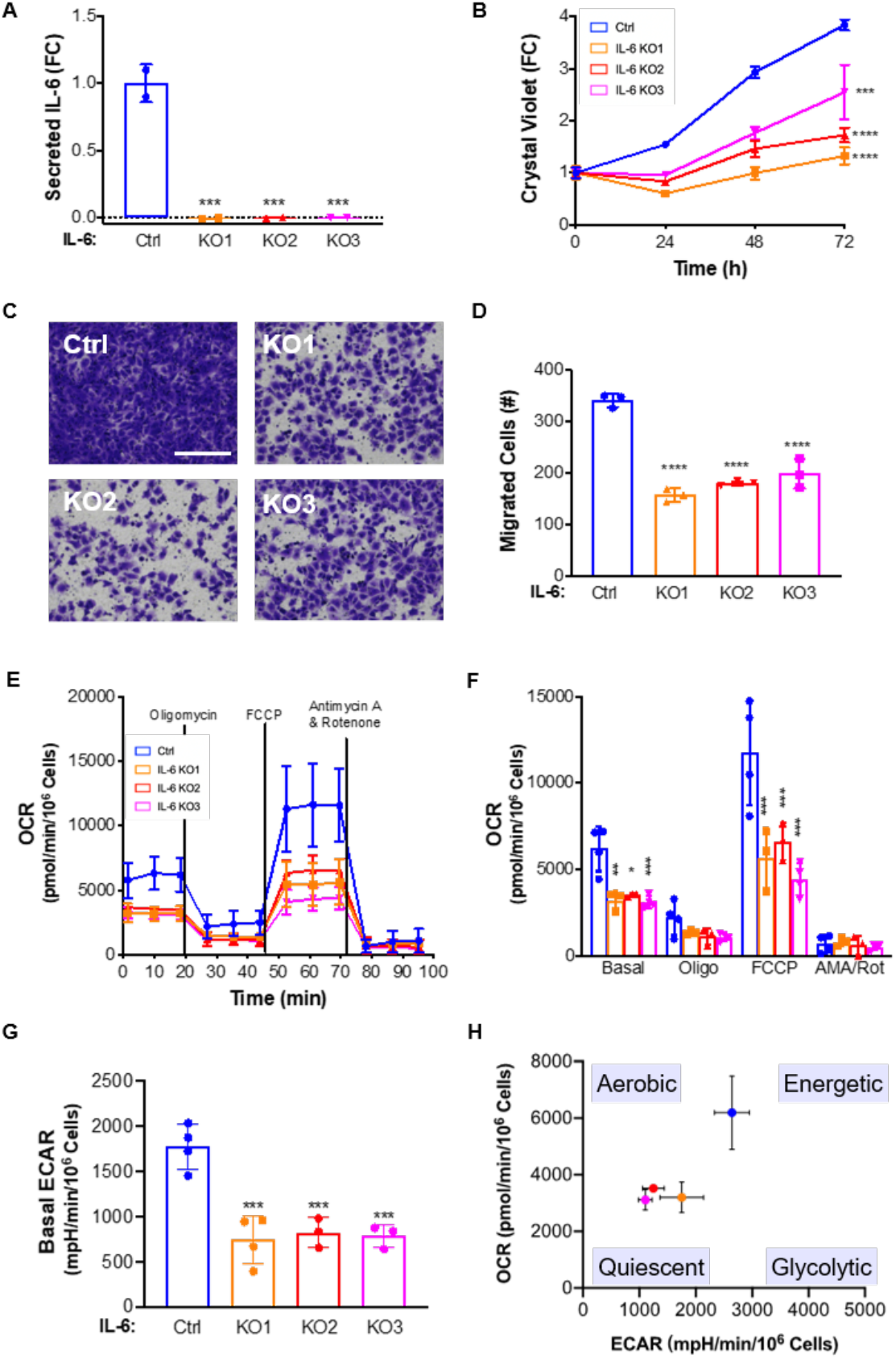
IL-6 knockout suppresses proliferation, migration, and induces a metabolic quiescent state in TSC2-deficient cells. (*A*) Secreted IL-6 is decreased in three IL-6 CRISPR/Cas9 clones compared to control, as measured by ELISA. (*B*) IL-6 knockout decreases the proliferation of TSC2-deficient MEFs compared to control as measured by crystal violet staining as a readout of cell density. (*C*) Representative images of (*D*) quantified cells migrated through transwells towards serum, which was decreased in IL-6 knockout, TSC2-deficient MEFs compared to TSC2-deficient control MEFs. Scale bar = 100um. (*E*) OCR is decreased in TSC2-deficient MEFs with IL-6 knockout compared to control cells. Data show measurements from the Seahorse extracellular flux analyzer using the MitoStress assay. (*F*) Summary and statistical analysis of OCR results. (*G*) Basal ECAR is decreased in TSC2-deficient MEFs with IL-6 knockout MEFs compared to control. mpH, milli-pH. (*H*) Energy map showing the global bioenergetic status of TSC2-deficient MEFs with IL-6 knockout compared to control. Data presented as the mean ± SD of three-four independent experiments. One-Way ANOVA and Student’s t test were used for statistical analysis. *p < 0.05, **p < 0.01, ***p < 0.001, ****p < 0.0001.

To elucidate the mechanisms through which IL-6 knockout inhibits the proliferation of TSC2-deficient cells, we examined oxidative phosphorylation and glycolysis using the Seahorse XF Analyzer. The MitoStress Test Assay utilizes the effects of various mitochondrial targeted compounds on oxygen consumption rate (OCR) and extracellular acidification rate (ECAR) as a readout of the cellular potential for oxidative phosphorylation (OXPHOS) and aerobic glycolysis, respectively. Prior studies have shown that TSC2-deficient cells upregulate glycolysis and OXPHOS to sustain the high bioenergetic and anabolic demands of mTORC1 hyperactivation (33-35). We found an overall decrease in OCR in the IL-6 knockout, TSC2-deficient cells compared to TSC2-deficient control cells expressing IL-6 (**Fig. 2*E***). Statistical analysis of the OCR data demonstrated that both basal and maximal respiration (FCCP-induced) were significantly reduced ∼50% (p<0.001) in all three IL-6 knockout clones (**Fig. 2*F***). We also found that IL-6 knockout suppressed aerobic glycolysis by ∼50% (p<0.001, **Fig. 2*G***). The reduction in both OCR and ECAR suggests that IL-6 knockout shifts TSC2-deficient cells to a bioenergetic quiescent state (**Fig. 2*H***). Acute treatment with recombinant IL-6 (200pg/ml, 24 hours) had no impact on the OCR or glycolytic profile of IL-6 KO cells (**Fig. S2*F* and *G***).

Collectively, these data demonstrate that IL-6 knockout has a significant impact on proliferation, migration, and metabolism of TSC2-deficient cells, suggesting that IL-6 may be a previously unappreciated mediator of metabolic reprogramming in TSC.

### IL-6 promotes *de novo* serine synthesis in TSC2-deficient cells

Since we observed an IL-6 dependent reduction in oxidative phosphorylation and glycolysis, we investigated the role of IL-6 in TSC2-deficient cell metabolism by performing targeted metabolomics, measuring ∼270 unique metabolites by liquid chromatography/mass spectrometry (LC/MS) (36). The IL-6 knockout clones showed a distinctive metabolic signature (**Fig. S3*A***). There was some variability between the three clones, as expected from the process of single cell cloning; however, principal component analysis (PCA) confirmed that the clones were more similar to one another than the TSC2-deficient control cell line (**Fig. S3*B***).

We next performed metabolite set enrichment analysis using MetaboAnalyst software by curating a single list of consistently differentially regulated metabolites from pairwise comparisons of the control cells with each of the IL-6 knockout clones. The most significantly impacted pathway (p<0.0003) with a false discovery rate (FDR) less than 5% was glycine and serine metabolism (**Fig. S3*C***). Glycolytic intermediates can be diverted from the TCA Cycle for utilization in *de novo* serine synthesis and the Pentose Phosphate Pathway (PPP). In tumors, upregulation of *de novo* serine metabolism and PPP supports nucleotide metabolism and redox homeostasis (37, 38). TSC2-deficient cells with IL-6 knockout have a >2-fold reduction in 3-phosphoserine (p<0.001, **Fig. S4*A***) and a ∼25% reduction in total serine levels (p<0.0001, **Fig. S4*B***). Glycine was not measured in our samples and cysteine was not changed by knocking out IL-6 (**Fig. S4*C***). We also discovered a ∼40% reduction in ribose 5-phosphate, a PPP intermediate, and ∼50% reduction in purines following IL-6 knockout in TSC2-deficient cells (**Fig. S4*D* and *E***).

While early metabolites of the TCA cycle (citrate, aconitate, and isocitrate) were increased by >2-fold (**Fig. S4*F*-*H***), metabolites downstream of α-ketoglutarate, which can be produced by glutaminolysis, were unaffected or decreased by IL-6 loss (**Fig. S4*I*-*M***). One possible explanation for this result is that decreased flux of glycolytic intermediates into PPP and *de novo* serine synthesis following IL-6 knockout increases flux into the TCA cycle. These data suggest that IL-6 plays significant role in the metabolism of TSC2-deficient cells, particularly glucose utilization.

To further investigate the impact of IL-6 on glucose metabolism, we used U-^13^C glucose tracing and measured labeled metabolites after 0, 1, 3 and 24 hours (39), focusing on the fractional enrichment of metabolites directly related to glucose metabolism (**Fig. 3*A***). All statistical analysis was performed at the final 24-hour time point shown in the bar graph below the time course data. The M+3 forms of serine (**Fig. 3*B***) and 3-phosphoserine (**Fig. 3*C***) were decreased in the cells with IL-6 knockout compared to controls with intact IL-6 by ∼50% (p<0.0001). Enrichment of M+3 phosphoglycerate was equal between the IL-6 knockout lines and TSC2-deficient controls, suggesting that IL-6 selectively effects shuttling of glycolytic intermediates into the *de novo* serine synthesis pathways (**Fig. 3*D***). IL-6 knockout decreased glucose-derived M+2 α-ketoglutarate by ∼50% (p<0.001, **Fig. 3E**) and increased glucose-derived M+2 glutamate by ∼10% (p<0.01, **Fig. 3*F***).

**Figure 3.**
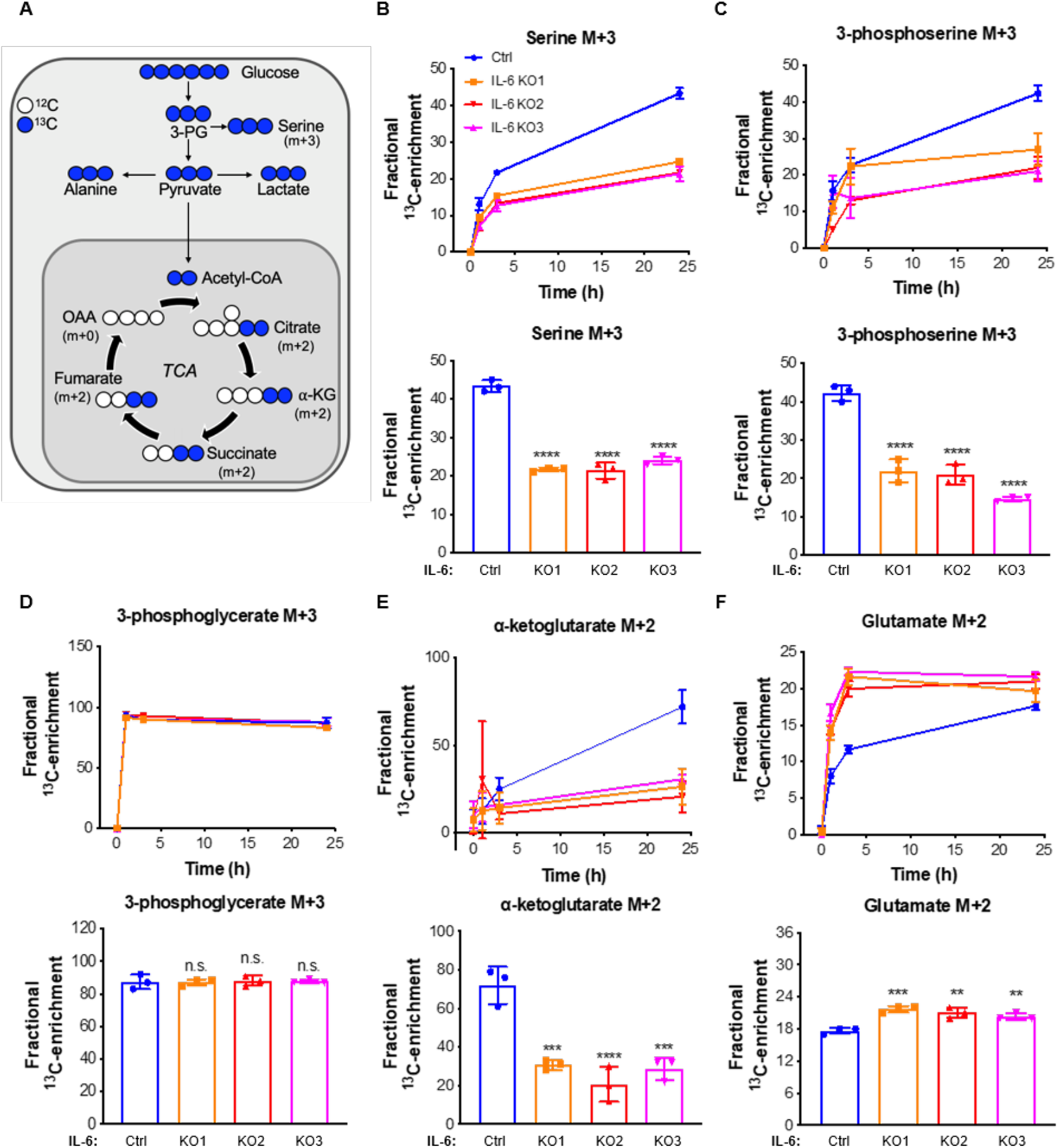
IL-6 regulates *de novo* serine synthesis in TSC2-deficient cells. (*A*) Schematic of U-^13^C-glucose metabolism. The incorporation of ^13^C atoms from ^13^C_6_-glucose into citrate, α-ketoglutarate (α-KG), succinate, fumarate, and oxaloacetate (OAA) are denoted as M+n, where n is the number of ^13^C atoms. (*B*-*F*). Fractional enrichment of the M+3 isotopologues of serine, 3-phosphoserine, and 3-phosphoglycerate or M+2 isotopologues of a-ketoglutarate and glutamate in TSC2-deficient MEFs with IL-6 knockout compared to TSC2-deficient controls (0, 1, 3, 24 hours after labelling; upper panels). Bar graphs show statistical analysis at the 24 h timepoint (bottom panels). Data presented as mean ± SD of three biological replicates. Statistical analysis performed by One-Way ANOVA, *p<0.05, **p<0.01, ***p<0.001, ****p <0.0001 as compared to control.

Glucose tracing data provide insights into the steady-state targeted metabolomics data, highlighting a significant reduction in the TCA cycle intermediate a-ketoglutarate derived from glucose. Furthermore, knocking out IL-6 in TSC2-deficient cells selectively suppressed glucose-derived intermediates from shuttling into *de novo* serine synthesis thereby impacting the production of serine.

### PSAT1 rescues proliferation of TSC2-deficient cells following IL-6 knockout

To identify the potential molecular mechanisms through which IL-6 regulates *de novo* serine synthesis in TSC2-deficient cells, we measured the mRNA expression of the key metabolic enzymes of the pathway, *PHGDH, PSAT1* and *PSPH* (**Fig. 4*A*-*D***). All three of the enzymes were significantly reduced at the mRNA level (>30%, p<0.01) in the IL-6 knockout clones compared to the control TSC2-deficient cells.

**Figure 4.**
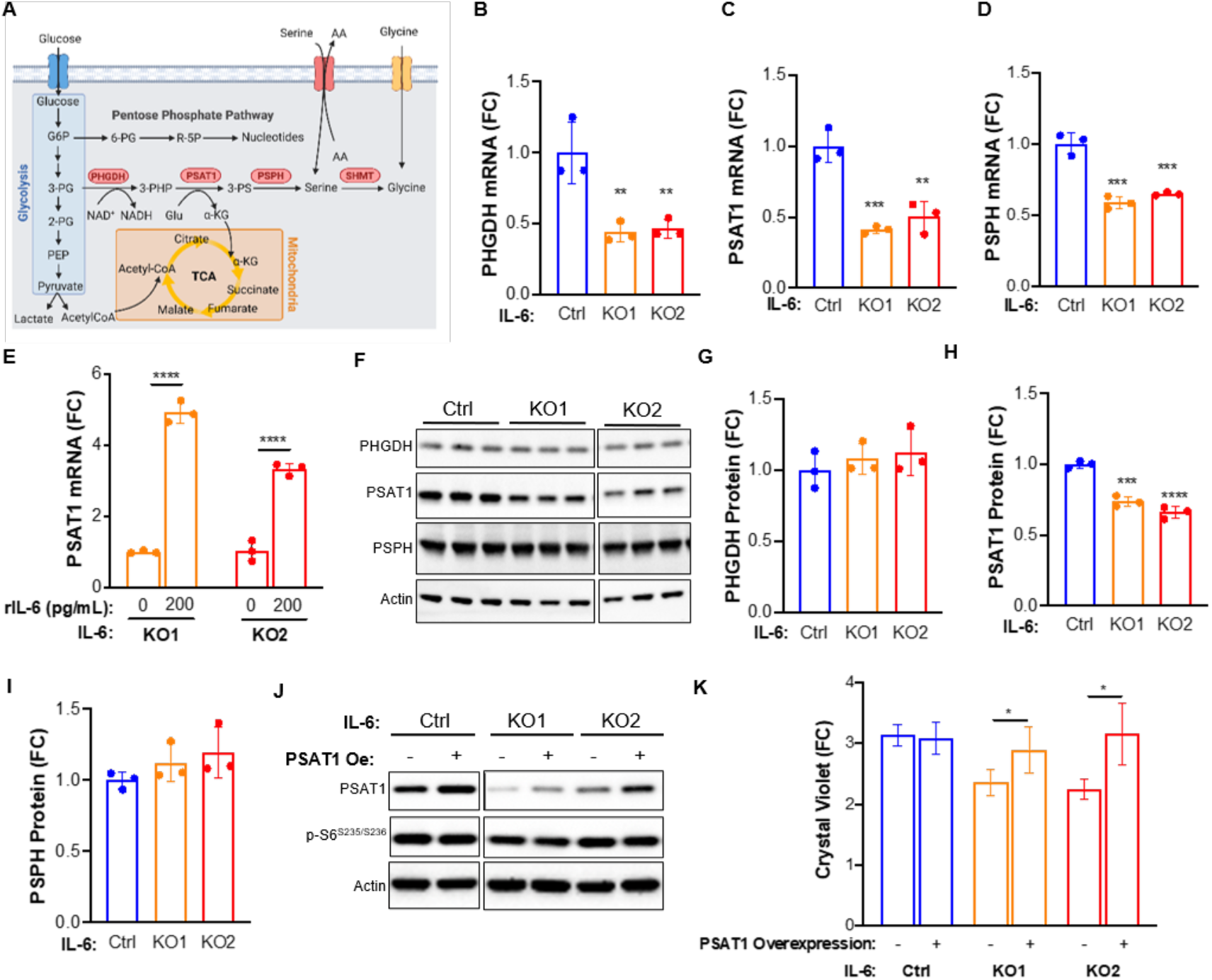
PSAT1 rescues proliferation of TSC2-deficient, IL-6 knockout cells. (*A*) Diagram depicting interrelationships between glycolysis, the pentose phosphate pathway (PPP), *de novo* serine biosynthesis, and the tricarboxylic acid cycle (TCA). (*B-D*) PHGDH, PSAT1 and PSPH mRNA levels are decreased in TSC2-deficient, IL-6 knockout cells compared to TSC2-deficient control MEFs. (*E*) PSAT1 mRNA levels in IL-6 knockout, TSC2-deficient cells are rescued upon recombinant IL6 (rIL-6) treatment (200 pg/ml; 24 h). (*F*-*I*) Western blot and densitometry showing expression of *de novo* serine biosynthesis enzymes and decreased PSAT1 expression in IL-6 knockout cells compared to control cells. Blots are from the same gel. Some lanes were cropped out for visualization purposes. (*J*) Western blot confirming PSAT1 overexpression in TSC2-deficient, IL-6 knockout cells and TSC2-deficient controls. Blots are from the same gel. (*K*) PSAT1 overexpression rescues proliferation of IL-6 knockout, TSC2-deficient cells (72 h; fold change relative to day 0, when the cells were washed and put into serum-free media). The data are presented as the mean ± SD of three independent experiments. One-Way ANOVA was used for statistical analysis. *p<0.05, **p<0.01, ****p <0.0001 as compared to control.

We next confirmed the dependency of *PSAT1* on IL-6 signaling in the TSC2-deficient cells using siRNA for IL-6Ra. A 90% reduction in IL-6Ra significantly reduced *PSAT1* expression ∼20% (p<0.01, **Fig. S5*A***). Interestingly, knocking down IL-6 ∼80% with siRNA had no effect on PSAT1 expression (**Fig. S5*B***), suggesting that TSC2-deficient cells are responding at least in part to binding of extracellular IL-6 to the IL6-Ra in a cell autonomous manner to regulate PSAT1 and *de novo* serine biosynthesis.

Importantly, treatment with recombinant IL-6 (rIL-6) rescued the expression of PSAT1 in the knockout clones, inducing PSAT1 expression by >3-fold (p<0.0001, **Fig. 5*E***). Recombinant IL-6 had no effect on *de novo* serine synthesis enzymes in the TSC2-wildtype MEFs (**Fig. S5*C***). Recombinant IL-6 also induced a >1.5-fold increase in PHGDH, PSPH and cytosolic SHMT1, but not mitochondrially localized SHMT2 (**Fig. S5*D* and *E***). These data highlight the regulation of *de novo* serine synthesis by IL-6 selectively in TSC2-deficient cells. IL-6 knockout reduced PSAT1 protein expression >25%, while PHGDH and PSPH protein expression were unchanged compared to the control TSC2-deficient MEFs (**Fig. 4*F*-*I***).

**Figure 5.**
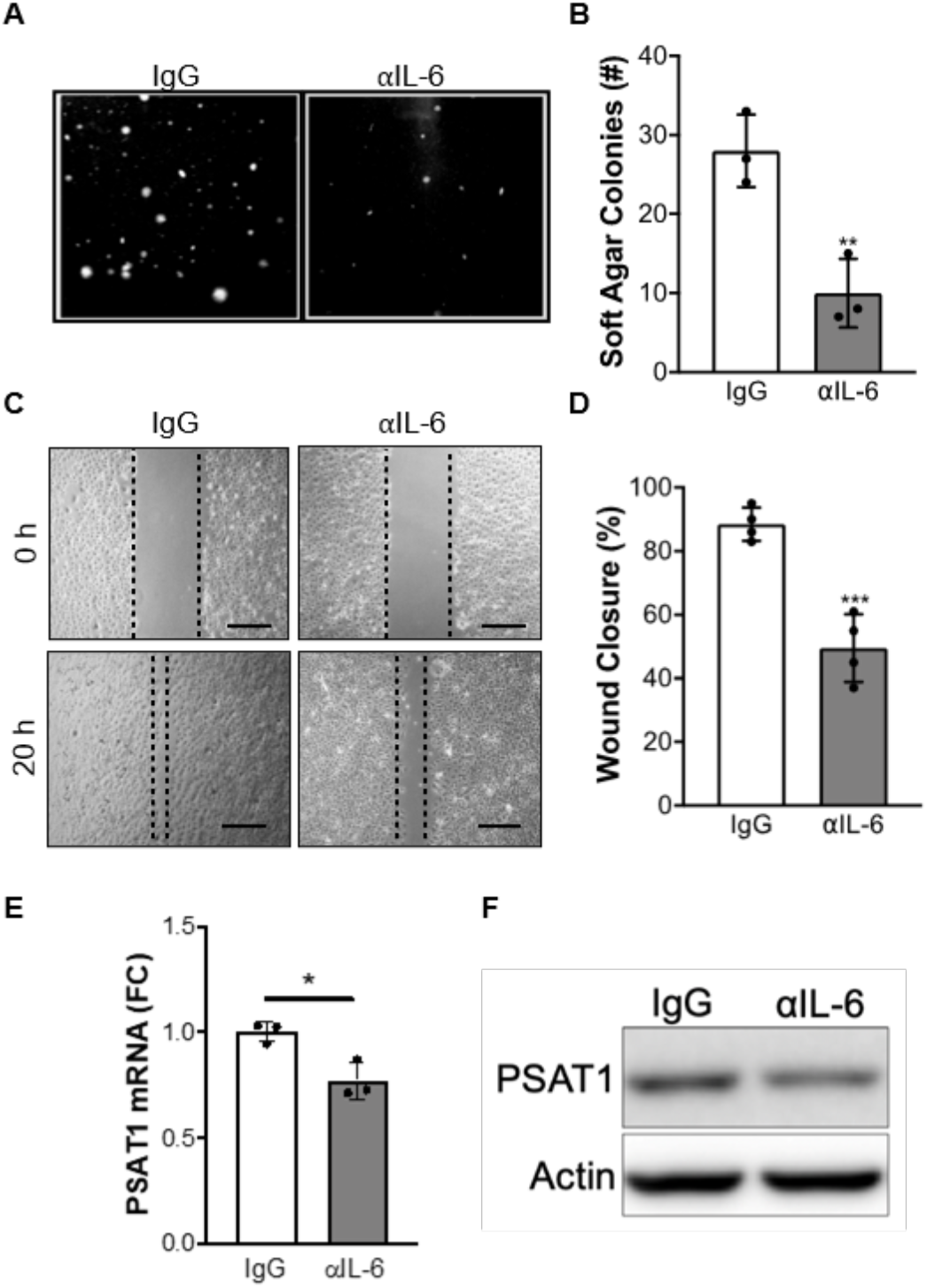
αIL-6 antibody suppresses proliferation, migration and PSAT1 expression in TSC2-deficient cells. (*A*) Representative images and (*B*) quantification of TSC2-deficient cell colonies grown for 14 days in soft agar and treated with αIL-6 or IgG control antibody (10ug/ml) in serum-free DMEM. Scale bar = 300μm. (*C*) Representative images and (*D*) quantification of wound-healing assay performed on TSC2-deficient cells treated with αIL-6 or IgG control antibody (10ug/ml). αIL-6 treatment decreases wound closure of TSC2-deficient cells compared to IgG control antibody after 20 h (αIL-6; 10ug/ml). Quantification of percent area filled between the two leading edges at 20 h compared to 0 h (0% filled). Scale bar = 150μm (*E*) PSAT1 mRNA and (*F*) PSAT1 protein are decreased by αIL-6 antibody (10ug/ml; 48 h) in TSC2-deficient cells. Cells were grown in serum-free DMEM for the duration of the experiments. The data are presented as mean ± SD of three independent experiments. Student’s t test was used for statistical analysis with *p<0.05, **p<0.01, ***p<0.001.

In order to gain insights into the mechanisms by which IL-6 may regulate the enzymes of *de novo* serine synthesis, we performed additional knockdown experiments. Using STAT3 siRNA and inducible Raptor and Rictor knockout MEFs (40), we determined that PSAT1 is regulated in a STAT3-*independent* and mTORC1-*dependent* manner (**Fig. S6*A*-*C***). These data are consistent with the finding that PSAT1 mRNA is increased ∼3-fold in TSC2-deficient cells compared to wildtype control cells (**Fig. S6*D* and *E***), as described in previous literature reports (41). In these prior publications, activating transcription factor 4 (ATF4), has been implicated in PSAT1 regulation downstream of mTORC1. We discovered a ∼15% decrease in *ATF4* mRNA by IL-6 knockout (p<0.001, **Fig. S6*F***). Interestingly, ATF4 protein expression and phosphorylation of S6 kinase were decreased in IL-6 knockout cells, compared to IL-6 expressing cells (**Fig. S6*G*)**. Importantly, *ATF4* expression was induced ∼2-fold (p<0.001) by rIL-6 treatment in IL-6 knockout TSC2-deficient MEFs compared to TSC2-wildtype MEFs, in which *ATF4* expression was unchanged by rIL-6 (**Fig. S6*H* and *I***). In order to determine the dependence of TSC2-deficient cells on PSAT1 downstream of IL-6, we overexpressed PSAT1 in the IL-6 knockout cells. Overexpression of PSAT1 was sufficient to rescue the proliferation of the IL-6 knockout cells and had no effect on the proliferation of the TSC2-deficient control cells (**Fig. 4*J* and *K***).

In summary, these data suggest that IL-6 promotes the proliferation of TSC2-deficient cells in a cell autonomous manner via the upregulation of PSAT1 and induction of *de novo* serine synthesis.

### IL-6 neutralizing antibody suppresses PSAT1 expression, proliferation, and migration in TSC2-deficient cells

IL-6 and IL-6Ra neutralizing antibodies are approved by the Food and Drug Administration of the United States for the treatment of various autoimmune diseases, including rheumatoid arthritis (42). Therefore, we sought to determine whether anti-IL-6 (aIL-6) antibody treatment, would impact the proliferation and migration of TSC2-deficient cells. We used p-STAT3^Y705^ expression as a surrogate marker of aIL-6 antibody activity (**Fig. S7*A* and *B***). In order to more closely mimic the long-term consequences of aIL-6 antibody on the TSC2-deficient cells we treated cells for 14 days and then quantified colony formation in anchorage-independent soft agar conditions. aIL-6 antibody reduced colony formation by ∼60% (p<0.01) compared to TSC2-deficent cells treated with control IgG antibody (**Fig. 5*A* and *B***). aIL-6 antibody also acutely reduced the migration of TSC2-deficient cells ∼40% compared to IgG antibody control (p<0.001, **Fig. 5*C* and *D***).

We next wanted to determine if aIL-6 antibody treatment inhibited proliferation and migration effects by limiting *de novo* serine synthesis, as we observed in the IL-6 knockout cells. We discovered that aIL-6 significantly reduced PSAT1 expression and reduced the M+3 labeling of serine in the TSC2-deficient cells (**Fig. 5 *E, F* and S7*C***). These data suggest that targeting IL-6 regulates PSAT1 expression and *de novo* serine synthesis in TSC2-deficient cells, and may be a therapeutic approach for the treatment of TSC and LAM.

### IL-6 neutralizing antibody suppresses the progression of renal tumors in TSC2^+/-^ mice

We next investigated the therapeutic impact of aIL-6 antibody in Tsc2^+/-^ mice, a well-established preclinical model of TSC, which spontaneously develops renal cysts and cystadenomas by 6 months of age (43). In the first set of experiments, we determined the therapeutic benefits of aIL-6 antibody as a single agent. Eight-month old Tsc2^+/-^ mice were treated with aIL-6 antibody or control antibody (IgG) (200 ug, 3 times a week) for one month (**Fig. 6*A***) and harvested 24-48h after the final injection. Kidneys were inspected macroscopically for gross cysts and tumor burden (**Fig. 6*B***). aIL-6 antibody reduced both the gross and microscopic tumor burden ∼30% (p<0.05) compared to the IgG treated mice (**Fig. 6*B*-*D***). aIL-6 antibody also reduced the number of proliferating Ki67+ cells in the renal lesions (30%, p<0.01, **Fig. 6*E* and *F***). Semi-quantitative analysis of PSAT1 expression in renal lesions of the Tsc2^+/-^ mice was performed by a blinded observer. Two out of three aIL-6 antibody treated mice showed a decrease in the PSAT1 expression across >10 lesions per mouse (**Fig. S8A and B**).

**Figure 6.**
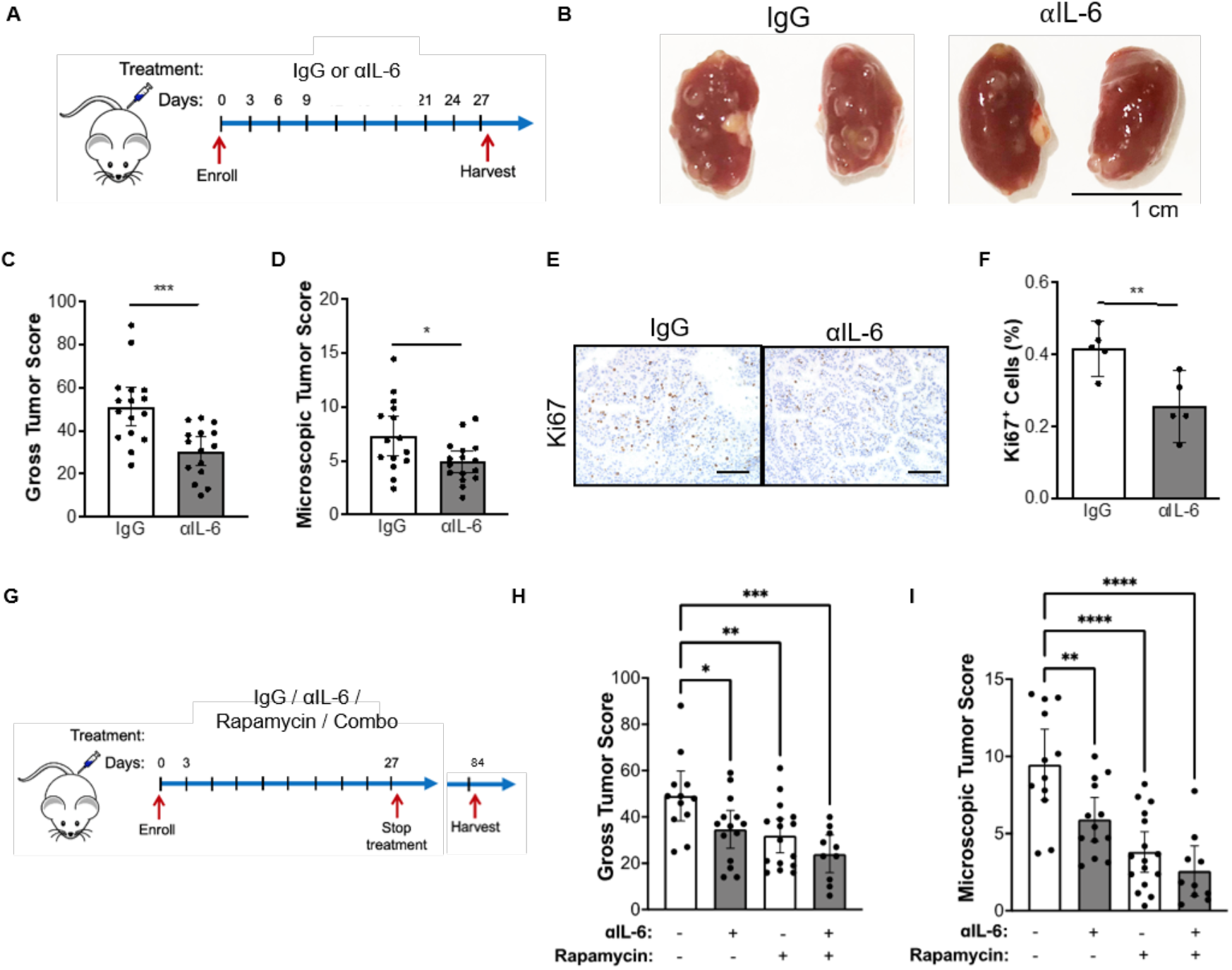
αIL-6 antibody suppresses renal cystadenoma formation in *Tsc2*^*+/-*^ mice. (*A*) Experimental design of αIL-6 antibody treatment. Tsc2+/-mice were injected intraperitoneally with IgG or αIL-6 antibody (200 ug/mouse, three times per week) for one month and then harvested 24-48h after the last injection. (*B*) Representative kidneys of *Tsc2*^*+/-*^ mice injected intraperitoneally with IgG or αIL-6 antibody. (*C*) αIL-6 antibody decreased the gross tumor score and (*D*) microscopic tumor score of *Tsc2*^*+/-*^ kidneys compared to IgG treated. (*E*) Representative images of Ki67 staining in Tsc2+/-renal tumors and (*F*) quantification showing decreased proliferation in mice treated with αIL-6 antibody compared to IgG control mice. (*G*) Experimental design to determine the duration of therapeutic benefit following αIL-6, rapamycin, or combination treatments. Tsc2+/-mice were treated for one month and then harvested 2 months after the final injection. (*H*) Gross tumor score and (*I*) microscopic tumor score of kidneys from *Tsc2*^*+/-*^ mice treated with IgG, αIL-6 (200ug/mouse, three times/week), rapamycin (3mg/kg three times/week) or combination, two months after treatment cessation. Data are presented as the mean +/-95% CI, each dot represents one kidney, Statistical analysis was performed with Student’s t test or One-Way ANOVA. Scale bar =100μm.

mTORC1 inhibition with rapamycin and related “rapalogs” is currently approved for patients with LAM and TSC. In LAM patients, lung function decline stabilizes, and many brain lesions and renal tumors shrink on mTORC1 inhibitor treatment in patients with TSC (44, 45). However, disease burden rapidly rebounds after treatment cessation, necessitating novel therapeutic interventions. To determine the lasting effectiveness of aIL-6 antibody treatment compared to rapamycin or combination therapy (**Fig. 6*G***), Tsc2^+/-^ mice at 5-6 months of age were randomly assigned to receive IgG, aIL-6 (200 ug, 3 times a week), rapamycin (3mg/kg 3 times a week) or the combination (aIL-6 and rapamycin) for one month. The mice were harvested 2 months after the final treatment. All three treatment arms showed significant reduction in tumor burden compared to the IgG control mice (**Fig. 6*H* and *I***). The mean gross tumor score was reduced by 25% by aIL-6 antibody (p<0.01), 30% by rapamycin (p<0.0001), and 50% by the combination (p<0.0001) compared to the IgG treated controls. The mean microscopic tumor burden was reduced by 40% by aIL-6 antibody (p<0.01), 60% by rapamycin (p<0.0001), and 70% by the combination (p<0.0001) compared to the IgG treated controls.

In summary, we observed a significant benefit from aIL-6 antibody treatment and long-term benefits of 1-month treatment with aIL-6 antibody. The combination of aIL-6 antibody and rapamycin appears to have an additive effect on tumorigenesis. These studies support the therapeutic potential of using clinically available approaches to target IL-6 in patients with TSC and LAM alone or in combination with mTORC1 inhibition.

## Discussion

We have discovered that IL-6 cooperates with mTORC1 activation to support TSC-associated metabolic reprogramming. Using steady state metabolomics in combination with U-^13^C glucose tracing we demonstrate that IL-6 plays a role in shunting glycolytic intermediates into *de novo* serine synthesis, a process which supports redox homeostasis and nucleotide metabolism in numerous tumors (41, 46-48). Our data suggest that IL-6 targeting has a significant impact on the metabolism and proliferation of TSC2-deficient cells. Interestingly, PSAT1 overexpression was sufficient to rescue the proliferation of the IL-6 knockout cells in the TSC2-deficient state, suggesting that regulation of serine metabolism is a key pathway downstream of IL-6. Serine, Glycine, and One Carbon (SGOC) metabolism fuels into numerous essential metabolic pathways (48, 49). Future studies will explore additional processes downstream of IL-6 and serine metabolism in the setting of mTORC1 hyperactivation. Of note, serine metabolism was shown to play a role in epigenetic processes following LKB1 loss and mTORC1 activation in KRAS-driven tumors (50). Serine availability is also important for the generation of certain lipid species, known to be dysregulated in TSC2-deficient cells, suggesting another aspect of TSC biology that may be significantly impacted by inhibiting IL-6 (47, 51). These studies highlight the far-reaching effects on features of TSC pathogenesis which may be mediated by IL-6 and serine metabolism.

IL-6 is a known autocrine, paracrine and endocrine factor that has been implicated in the initiation, progression and metastasis of numerous tumor types including skin, breast, lung, and kidney (13, 52, 53). In this study we focused on the cell autonomous roles of IL-6 on the metabolism and tumorigenesis of TSC2-deficient cells. Interestingly, we discovered that circulating IL-6 levels are elevated in the serum of LAM patients, and in TSC2-deficient human angiomyolipoma cells. Additionally, expression of IL-6Rα is elevated in TSC2-deficient cells suggesting that IL-6 plays a role in a cell autonomous manner. Interestingly, IL-6Rα can be shed into the extracellular milieu, allowing previously non-responsive cells expressing the ubiquitous gp130 receptor to sense IL-6 (42, 54, 55). This mechanism of action highlights the exciting possibility that TSC2-deficient cells may also shed IL-6Rα to modulate the tumor microenvironment, exerting non-cell autonomous effects. Since recent studies have shown that TSC-associated pulmonary LAM and renal angiomyolipomas are responsive to immune checkpoint blockade (28, 56), the importance of the immune system in TSC and LAM disease progression has been well established. Importantly, IL-6 targeted therapies following mTORC1 activation caused by STK11/LKB1 loss in RAS-driven lung tumors decreased the immunosuppressive effects of tumor associated neutrophils (57). Therefore, understanding the impact of IL-6 mediated signaling on the tumor microenvironment in TSC and LAM will be a critical next step for future studies.

IL-6 activates the pro-oncogenic transcription factor STAT3 via binding to IL-6Ra and gp130, leading to canonical JAK/STAT signaling. STAT3 activation is a well-described feature of TSC (15, 16, 19, 21, 23, 58, 59), including both brain and lung manifestations (16, 20). However, the mechanisms underlying this activation are not completely understood. We propose a model in which secreted IL-6 potentiates the STAT3 signal in TSC2-deficient cells. STAT3 is also directly activated by mTORC1-mediated phosphorylation on serine 727, a phosphorylation event required for maximal transcriptional activation (60). Together these data support the activation of a STAT3/IL-6 positive feedback loop in TSC (61). Surprisingly, we discovered that IL-6 regulates *de novo* serine synthesis in a STAT3-independent manner in TSC2-deficient cells. Interestingly, the transcription factor, ATF4, known to regulate *de novo* serine synthesis enzymes (41, 62), was both mTORC1-dependent and regulated by IL-6. IL-6 activates additional pro-tumorigenic pathways including RAS, phosphatidyl inositol 3-phosphate (PI3K-AKT), and Yes associated protein/taffazin (YAP/TAZ) (13, 63-65). Furthermore, the IL-6 gene promoter contains an antioxidant response element and can be regulated by Nuclear Factor Erythroid 2 (NRF2) (66), which is upregulated in TSC lesions (67), and has been previously implicated in regulating *de novo* serine synthesis in tumors (68). Finally, YAP/TAZ signaling has been shown to be upregulated in TSC and plays a role in the regulation of transaminases in cancers (69, 70). Our data suggest that the regulation of *de novo* serine synthesis downstream of IL-6 may involve a complex interplay of transcription factors, many of which are implicated in TSC and LAM pathogenesis.

This project highlights potential new therapeutic strategies for TSC and LAM. To assess the therapeutic potential of targeting IL-6 in TSC and LAM, we treated 8-month-old *Tsc2*^*+/-*^ mice that have established tumor burden with IL-6 neutralizing antibody (αIL-6 antibody). We demonstrate that αIL-6 antibody suppresses the formation of renal cysts and cystadenomas in *Tsc2*^*+/-*^ mice. IL-6 targeted therapies are currently FDA-approved for the treatment of rheumatoid arthritis and Castleman’s Disease (54, 55). Interestingly, we found only a partial reduction of IL-6 upon rapamycin treatment *in vitro* and an additive benefit of combining aIL-6 antibody with rapamycin *in vivo*. These data suggest that IL-6 pathway targeted therapies may work well in combination with mTORC1 inhibition, which is standard of care for most patients with progressive TSC and LAM. Efforts are also underway to develop novel inhibitors of serine metabolism. These studies have shown that tumors must be in a serine limited environment for maximal therapeutic potential. The brain is one organ system with low serine and glycine in which PHGDH inhibitors have shown therapeutic benefit in preclinical models (71), with possible implications for the various brain tumors that form in TSC. Finally, a serine/glycine limited diet has also been shown to improve the efficacy of these inhibitors in tumor models (72, 73).

In summary, we have uncovered a distinct IL-6 dependent metabolic signature, which plays an important role in supporting the proliferative and bioenergetic activity of TSC2-deficient cells. In particular, IL-6 appears to promote *de novo* serine biosynthesis and expression of the key intermediate enzyme, PSAT1. Targeting IL-6 with a neutralizing antibody exerts acute and durable therapeutic responses alone and in combination with rapamycin in vivo. Future studies may elucidate the molecular mechanisms by which IL-6 regulates *de novo* serine biosynthesis and the potential therapeutic benefits of directly targeting *de novo* serine metabolism in TSC and LAM.

## Materials and Methods

### Cell lines and treatment

*Tsc2*^*-/-*^*p53*^*-/-*^ and *Tsc2*^*+/+*^*p53*^*-/-*^ mouse embryonic fibroblasts (MEF) were provided by David Kwiatkowski (Brigham and Women’s Hospital, Boston, MA). The MEFs and stocks prepared at passage 8. TTJ cells and 105K were derived from a renal cystadenoma of a C57BL/6 *Tsc2*^*+/–*^ mouse, with re-expressed Tsc2 or empty vector, as previously described (28). The 621-101 cells were isolated from a patient renal angiomyolipoma and immortalized with E6/E7, and the parental 621-101 cells were then use to re-express Tsc2 (621-103) or empty vector (621-102), as previously described (74, 75). HEK293 cells were obtained from ATCC. The inducible raptor and rictor knockout MEFs were generated in the laboratory of Michael Hall, the cells were cultured in 10% FBS in DMEM and treated with 1uM of tamoxifen or ethanol control for 72 h to induce knockout prior to harvest (40). For IL-6 knockout, TSC2-deficient cells were transduced with single lentivirus containing an spCas9 and sgRNA (CTTCCCTACTTCACAAGTC) expression cassette to target spCas9 cleavage to IL-6. The lentiviral plasmids (LV01) and lentivirus production were obtained from Sigma-Aldrich. Cells were sorted for GFP expression by flow cytometry and maintained in puromycin (3μg/mL). IL-6 knockout was confirmed by Sanger sequencing and ELISA. For PSAT1 overexpression, lentiviral vector of PSAT1 (EX-Mm13089-Lv197-GS, GeneCopoeia) was transfected into HEK-293T cells along with lentiviral packaging mix to produce lentivirus. Tsc2-deficient cells with or without IL-6 overexpression were transfected with lentivirus and selected with blasticidin at 10μg/ml. Knockdown experiments were performed using Silencer Select siRNA from Ambion STAT3 (4392420) and IL-6RAa (4390771) transfected using Lipofectamine RNAiMax Reagent (Invitrogen). All cells tested negative for mycoplasma contamination using MycoAlert (Lonza) and were re-tested monthly. Cells were cultured at 37°C in 5% CO2 in DMEM supplemented with 10% FBS and gentamycin sulfate (50μg/mL). For serum-free conditions, cells were cultured in DMEM without serum.

### Antibodies and drugs

The following antibodies were used: TSC2 (Cell Signaling Technology, 4308S), S6 (2217S), pS6(2211L), IL-6 (Santa Cruz Biotechnology, sc-57315), STAT3 (9139S), pSTAT3 (9145S), PSAT1 (Protein Tech, #20180-1-AP), PHGDH (Protein Tech, 14719-1-AP), PSPH (Protein Tech, 14513-1-AP), β-actin (Sigma-Aldrich), Ki-67 (ebioscience, 14-5698-82). For immunohistochemistry, PSAT1 antibody (2102) was purchased form Origene. IL-6 neutralizing antibody and IgG control (Bioxcell, BE0046 and BE0088). Rapamycin and Torin1 were purchased from LC Laboratories. Recombinant mouse IL-6 (406-ML) was purchased from R&D.

### Seahorse assay

The MitoStress Test Assay and the Seahorse XFe24 analyzer were used. Cells were seeded into the XFe24 microplate and incubated for 24 hours in 10% FBS DMEM. The next day, cells were washed by PBS and cultured in FBS free DMEM for 24 hours. Add compounds (final concentrations: 1μM oligomycin, 1μM FCCP, 0.5μM rotenone/antimycin A) to pre-hydrated sensor cartridge. The sensor cartridge and XFe 24 microplate were placed in the XFe Analyzer. Results were normalized to cell number.

### Cytokine Array

*A* RayBiotech Human Cytokine Array 5 was used to detect differential secretion of cytokines from angiomyolipoma-derived 621-101 cells compared to human embryonic kidney HEK293 cells according to the manufacturer’s instructions. Cells were grown in IIA Complete Media and serum starved overnight (16h) in 3ml of media on 10cm dishes. One mL of media was applied to the cytokine array and incubated for 2 hours. As a control, one array was incubated with unconditioned media.

### ELISA

ELISA was performed using conditioned media (∼80% confluent cells, concentrated with Millipore UFC 900324 filters) and normalized by protein concentration (Bio-Rad Laboratories, #5000006). Levels of secreted IL-6 were determined according to the manufacturer’s protocol (R&D Systems, IL-6 Quantikine ELISA Kit).

### Crystal violet assay

Cells were seeded at a density of 1,000 cells/well in 96-well plates and changed into serum-free media after 24 hours. At the indicated time points, cells were fixed for 15 minutes with 10% formalin and then stained with 0.5% crystal violet in distilled water for 20 minutes. Crystal violet was removed and cells were washed with water followed by drying at room temperature. Crystal violet was solubilized with 200 ml of methanol and measured with a plate reader (OD 540; BioTek, Winooski, VT, USA). Proliferation was assessed by comparing the change in OD 540 at 24, 48 and 72 h as normalized to 0h (start of serum-free proliferation) for each cell line.

### Transwell migration assay

Migration was evaluated as described previously (76). Briefly 5×10^4^ cells were seeded in 100 μl of serum-free DMEM in the upper chamber of a 6.5 mm polycarbonate Transwell with 8.0 μm pores (Corning, USA), through which the cells were allowed to migrate for 6 h at 37 °C toward 10% FBS in the basal compartment. At the end of the incubation, migrated cells on the lower surface of the transwell were fixed, stained, and quantified.

### Soft agar assay

Soft agar assays were performed to measure anchorage-independent growth. Briefly, 5 × 10^3^ cells were placed into a single well in a 6-well plate. Cells were embedded into 0.4% low-melting agarose (Sigma) and layered on top of a 0.8% agarose base. After 2 weeks of growth, the cells were fixed and analyzed. Colony number was quantitated using ImageJ (v1.53).

### Wound healing assay

Cells were seeded into six-well plates in DMEM culture medium and allowed to grow for 24 hours until confluency was reached. Cells were then washed with 1x PBS, and a scratch was made using a 200 μl tip at the center of the well. The monolayers were imaged at the indicated times using a light microscope at 100x magnification. The results were quantified using ImageJ (v1.53) software.

### RNA extraction and quantitative real-time PCR

Total RNA was extracted using RNeasy Mini Kit (QIAGEN, USA). The RNA concentration was measured using a Nanodrop 2000c (Thermo Scientific, USA). Two micrograms of RNA were reverse transcribed using a High-Capacity cDNA Reverse Transcription Kit (Thermofisher, USA) with random primers. For qPCR, a Taqman-based method was used and the relative quantitation of gene expression was determined using the comparative CT (ΔΔCT) method and normalized to β-actin gene and a calibrator sample that was run on the same plate. PCR primers and probe sets were obtained from Thermofisher, USA: IL-6 (assay ID Mm00446191_m1, 124 bp amplicon length), PSAT1 (assay ID Mm04932904_m1, 109 bp amplicon length), and β-actin control (cat# 4351315).

### Western blot analysis

After indicated treatment, live cells were lysed on ice in 1× RIPA (Cell Signaling Technology) containing phosphatase and protease inhibitors. For the membrane fractionation experiments Mem-PER™ Plus Membrane Protein Extraction Kit was used according to manufacturer’s instructions (Thermo Scientific, 89842). The concentration of proteins was determined using a Bio-Rad Protein Assay Dye Reagent Concentrate (Bio-Rad Laboratories, #5000006). A total of 15 μg of protein from each sample were mixed with NuPAGE™ LDS Sample Buffer (Thermofisher, NP0007) and Reducing Sample Buffer (Invitrogen, NP009, USA), resolved on a 4-12% Bis-Tris gels (Thermofisher), then transferred to PVDF membranes (MilliporeSigma, USA). Blots were blocked with 5% milk and incubated with primary and second antibodies. Chemiluminescence was captured with Syngene G-Box gel documentation system.

### Immunohistochemistry staining

Kidneys were formalin-fixed, paraffin-embedded, and tissue was cut in 3-to 4-μm sections then air-dried overnight. The sections were deparaffinized, rehydrated, and subjected to heat-induced epitope retrieval using low pH target retrieval solution for 15 minutes. Sections were incubated with Ki67 of PSAT1 primary antibody (1:100 dilution). Slides were developed using DAB and counterstained with hematoxylin.

### Targeted Mass Spectrometry

Samples were re-suspended using 20 mL HPLC grade water for mass spectrometry. 5-7 μL were injected and analyzed using a hybrid 6500 QTRAP triple quadrupole mass spectrometer (AB/SCIEX) coupled to a Prominence UFLC HPLC system (Shimadzu) via selected reaction monitoring (SRM) of a total of 270 endogenous water-soluble metabolites for steady-state analyses of samples. Some metabolites were targeted in both positive and negative ion mode for a total of 305 SRM transitions using positive/negative ion polarity switching. ESI voltage was +4950V in positive ion mode and –4500V in negative ion mode. The dwell time was 3 ms per SRM transition and the total cycle time was 1.39 seconds. Approximately 10-14 data points were acquired per detected metabolite. Samples were delivered to the mass spectrometer via hydrophilic interaction chromatography (HILIC) using a 4.6 mm i.d x 10 cm Amide XBridge column (Waters) at 400 μL/min. Gradients were run starting from 85% buffer B (HPLC grade acetonitrile) to 42% B from 0-5 minutes; 42% B to 0% B from 5-16 minutes; 0% B was held from 16-24 minutes; 0% B to 85% B from 24-25 minutes; 85% B was held for 7 minutes to re-equilibrate the column. Buffer A was comprised of 20 mM ammonium hydroxide/20 mM ammonium acetate (pH=9.0) in 95:5 water:acetonitrile. Peak areas from the total ion current for each metabolite SRM transition were integrated using MultiQuant v3.0 software (AB/SCIEX) (36, 39). Analysis of metabolomics data including the generation of heatmaps and metabolite set enrichment analysis (MSEA) was performed using the open access MetaboAnalyst Software (v4.0 or 5.0).

### U-^13^C Glucose Tracing

Cells (4×10^5^) were seeded onto 60mm plates in 10% FBS DMEM (ThermoFisher Scientific, Gibco #11995123). The next day the cells were washed with PBS and transferred to serum-free DMEM (Thermo Fisher Scientific, Gibco #11966025) supplemented with 4.5 g/L D-glucose. For U-^13^C Glucose Tracing following aIL-6 antibody experiments, the cells also were washed with PBS and transferred to either aIL-6 antibody (10 ug/ml) or IgG (10 ug/ml) in serum-free DMEM (Thermo Fisher Scientific, Gibco #11966025) supplemented with 4.5 g/L D-glucose. At −24h, −3h, −1h to harvest, the cells were washed with glucose-free and serum-free DMEM (Gibco #11966025) and then incubated in DMEM (#11966-025) supplemented with 4.5 g/L U-^13^C Glucose (Sigma Aldrich) for 0h, 1h, 3h, or 24h. The cells were harvested and analyzed as described above (36, 39).

### Human plasma specimens

Patient samples and healthy control samples were obtained through a Partner’s Health Care Institutional Review Board approved protocol. Subjects were consented by the clinical research team free of coercion and samples were collected, deidentified, and coded. Plasma Interleukin-6 was measured by ELISA (R&D Quantikine Human IL-6 ELISA or High Sensitivity IL-6 ELISA).

### Animal studies

All animal studies were performed in accordance with institutional protocols approved by BWH Institutional Animal Care and Use Committee. For the cystic kidney model, we crossed the CAGGCRE-ER^TM+/-^ (The Jackson Laboratory) to Tsc2 ^flox/flox^ (Michael Gambello). Recombination of Tsc2 was induced in 8-10 week old CAGGCRE-ER^TM+/-^; Tsc2 ^flox/flox^ or controls (CAGGCRE-ER^TM-/-^; Tsc2 ^flox/flox^) mice with intraperitoneal tamoxifen (in corn oil) at a dose of 1mg per day for 5 consecutive days. Kidneys were harvested at 5 months of age. *Tsc2*^*+/–*^ mice in the A/J background were generated in house as described previously (43, 77). Intraperitoneal injection was done with 200 ug of αIL-6 antibody or control rat IgG antibody (Bioxcell) three times per week as previously described (57). After a total of 4 weeks of treatment, mice were harvested and the severity of renal lesions was scored using previously established macroscopic and microscopic scoring methods (43, 77). Macroscopic cysts per kidney were scored according to size: < 1mm, score 1; 1–1.5, score 2; 1.5–2, score 5; and > 2, score 10. The sum of the cyst scores were determined and reported per kidney. Microscopic kidney tumor scores were determined by an observer blinded to the experimental conditions using a semi-quantitative algorithm and hematoxylin and eosin (H&E) sections of samples prepared by embedding 1 mm-interval sections. Each tumor or cyst identified was measured (length, width) and percent of the lumen filled by tumor determined (0% for a simple cyst, and 100% for a completely filled, solid tumor). The measurements were converted into a score using a previously established formula (43). Semi-quantitative analysis of PSAT1 immunohistochemistry was performed by a blinded observer who scored the PSAT1 signal on a sale from 0-5 (0-no signal, 5-maximum observed signal) in ∼10 renal lesions per mouse. The average score for each mouse was then calculated and the Log2 fold change of the aIL-6 antibody treated mice was calculated relative to the IgG control mice.

### Statistical analyses

Normally distributed data were analyzed for statistical significance with Student’s unpaired t-test and multiple comparisons were made with One-Way and Two-Way ANOVAs with Bonferroni correction. *In vivo* data are presented as the mean +/− 95% confidence interval (CI) and *in vitro* studies are presented as the mean +/− standard deviation (SD). Analysis was performed using GraphPad Prism version 8; GraphPad Software, www.graphpad.com. Statistical significance was defined as p < 0.05.

## Acknowledgments

This work was funded by the NIH (K01-DK116819 to HCL, R01HL146541 to WS, U01 HL131022-04 to EPH,) and The Engles Family TSC/LAM Research Fund. The mass spectrometry work was partially funded by NIH grants 5P01CA120964 (J.M.A.) and 5P30CA006516 (J.M.A.). We thank Dana-Farber/Harvard Cancer Center in Boston, MA for the use of the Rodent Histopahtology Core, which provided tissue embedding and sectioning services. Dana-Farber/Harvard Cancer Center is supported in part by a NCI Cancer Center Support Grant # NIH 5P30 CA06516. We would also like to acknowledge Clemens K. Probst for technical assistance with experiments. Schematics were generated using Biorender.com. This work was performed in part to meet the requirements of the doctoral thesis of Dr. Ji Wang from Zhejiang University School of Medicine, Hangzhou, China.

---

**Fig. S1.**
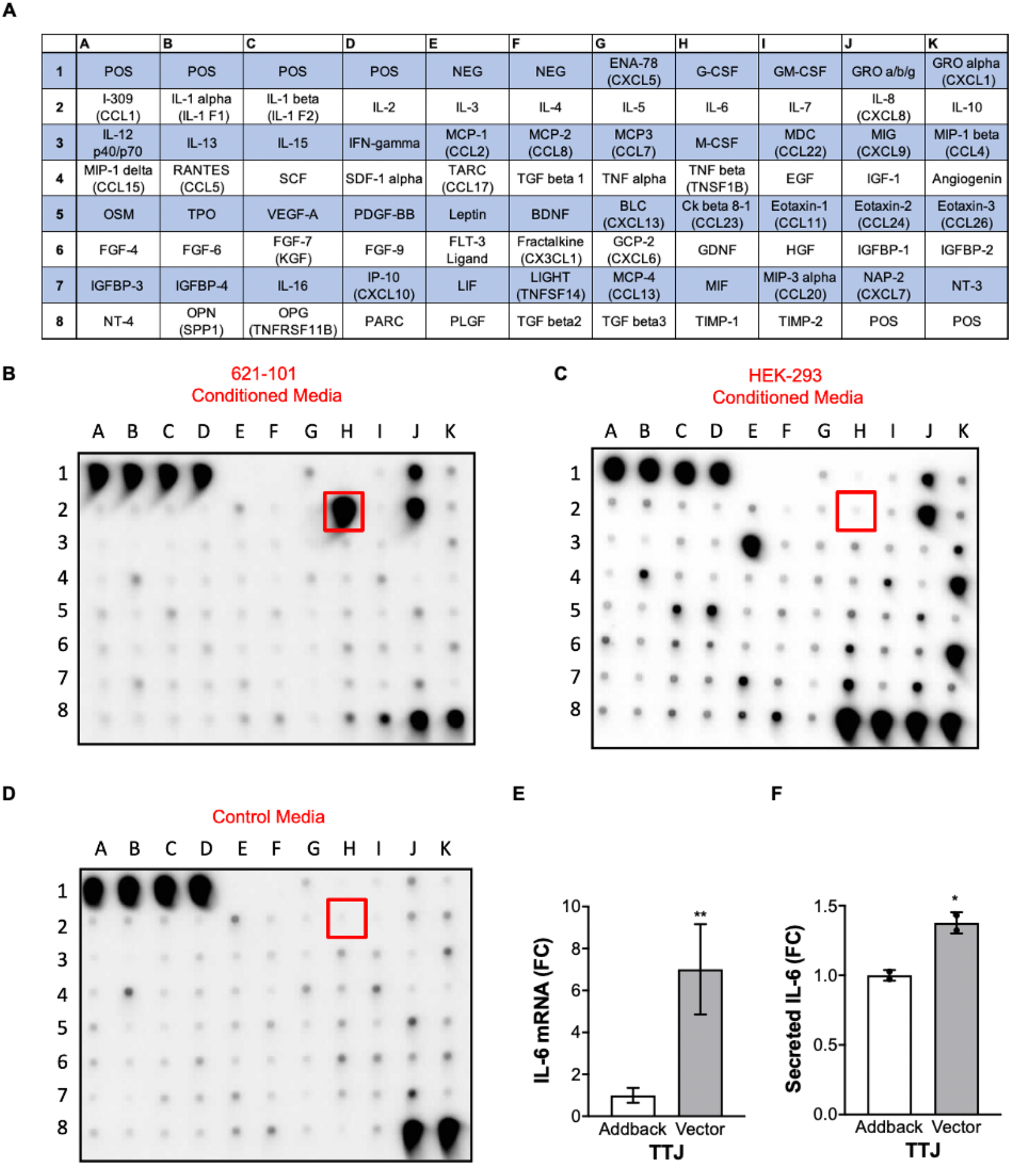
Increased IL-6 secretion in conditioned media from TSC2-deficient cells. (*A*) Layout of cytokine positions in RayBiotech Human Cytokine Array 5. (*B*) Cytokine array showing IL-6 expression is elevated in conditioned media from TSC2-deficient, patient-derived angiomyolipoma cells (621-101). (*C*) Cytokine array showing IL-6 expression in conditioned media from human embryonic kidney HEK293 cells. (*D*) Media incubated without cells was used as a blank control for the cytokine profile. (*E*) *IL-6* mRNA expression is increased in TSC2-deficient TTJ cells compared to TSC2 addback cells. (*F*) Secreted IL-6 is increased in conditioned media from TSC2-deficient TTJ cells compared to TSC2 addback cells.

**Fig. S2.**
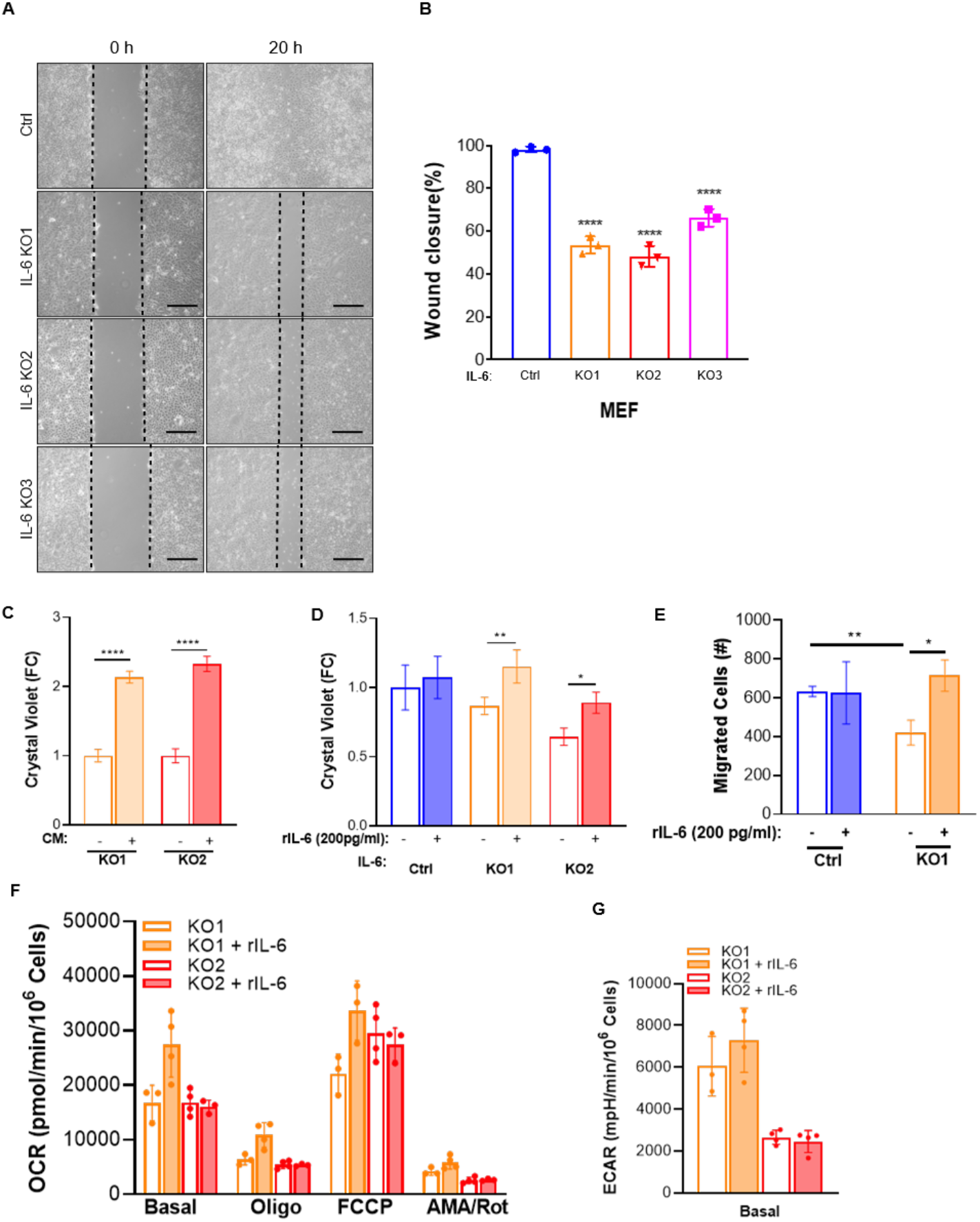
Rescue of IL-6 knockout cell proliferation, migration and bioenergetic assays. (*A*) IL-6 knockout using three separate CRISPR/Cas9 clones shows decreased wound closure compared to control cells. (*B*) Quantification of percent area filled between the two leading edges at 20 h compared to 0 h (0% filled). (C) The proliferation at 96h of IL-6 knockout, TSC2 deficient cells was rescued by conditioned media, generated by concentrating serum free media from TSC2 deficient cells at 24 hours. (D) The proliferation at 72h of IL-6 knockout, TSC2 deficient cells was rescued by rIL-6 (200pg/ml) treatment in serum free conditions. (E) rIL-6 rescued transwell migration of IL-6 knockout cells. Cells were pre-treated with rIL-6 (24h, 200pg/ml) and then seeded into the apical side of the transwell with rIL-6 or not treated controls. *F*) OCR and (*E*) ECAR are unchanged in IL6 KO cells upon treatment with recombinant IL6. Data show measurements from the Seahorse extracellular flux analyzer using the MitoStress assay. The data are presented as mean ± SD of three independent experiments. Students t test or One-Way ANOVA were used for statistical analysis.*\*p*<0.05, *\*\*p*<0.01, \*\*\*\**p* < 0.0001.

**Fig. S3.**
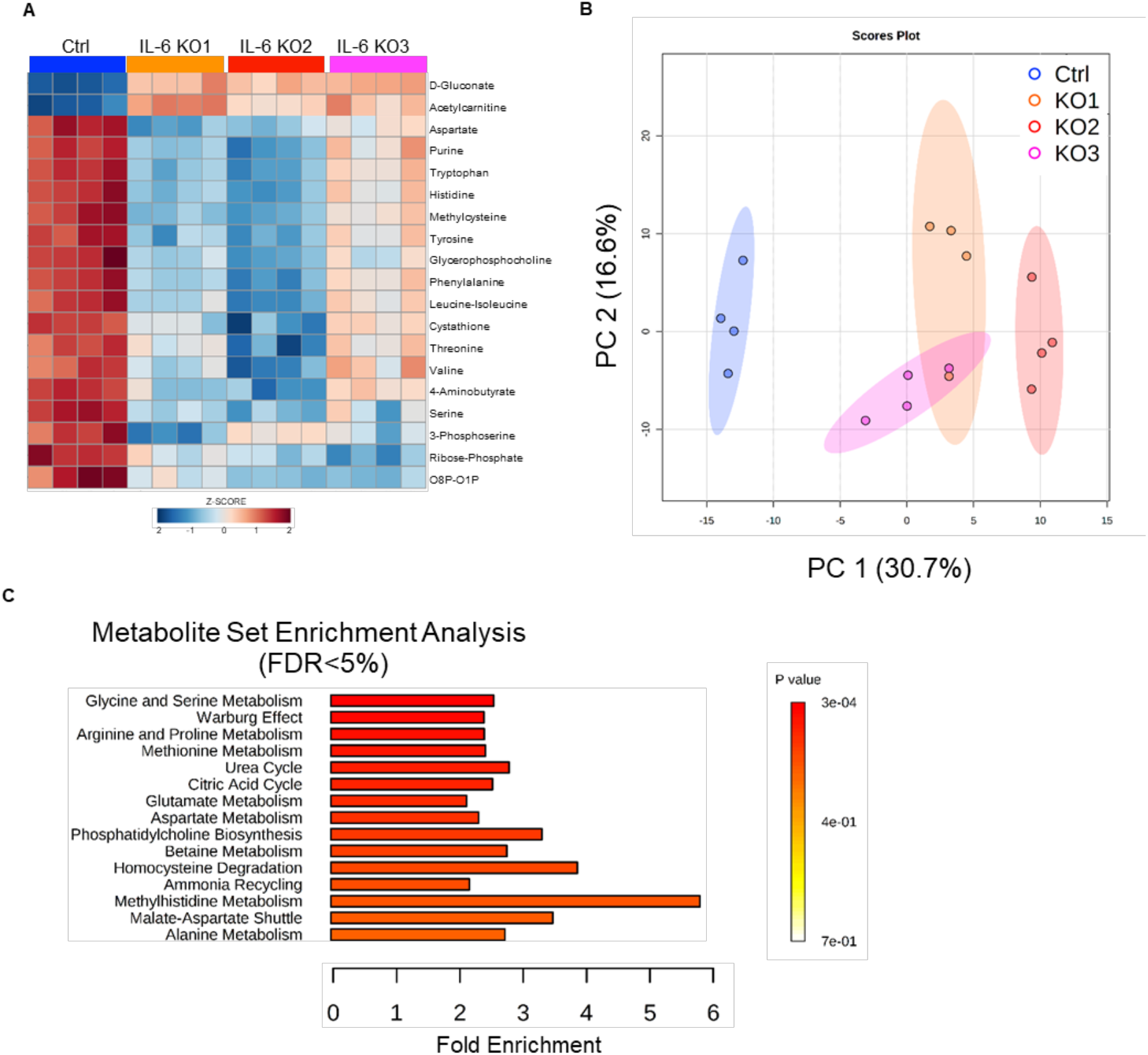
IL-6 knockout shows a distinct metabolic signature. (*A*) Hierarchical clustering and heatmap showing the top 20 differential metabolites in three IL-6 knockout TSC2-deficient MEF clones compared to control. (*B*) Principal component analysis shows that the metabolism of the three IL-6 knockout clones is distinctive from control cells. (*C*) Metabolite Set Enrichment Analysis (MSEA) of differentially regulated metabolic pathways upon IL-6 knockout with false discovery rate (FDR) <5% identifies Glycine and Serine Metabolism as the most significantly regulated pathway. Heatmaps and MSEA were generated using MetaboAnalyst.

**Fig. S4.**
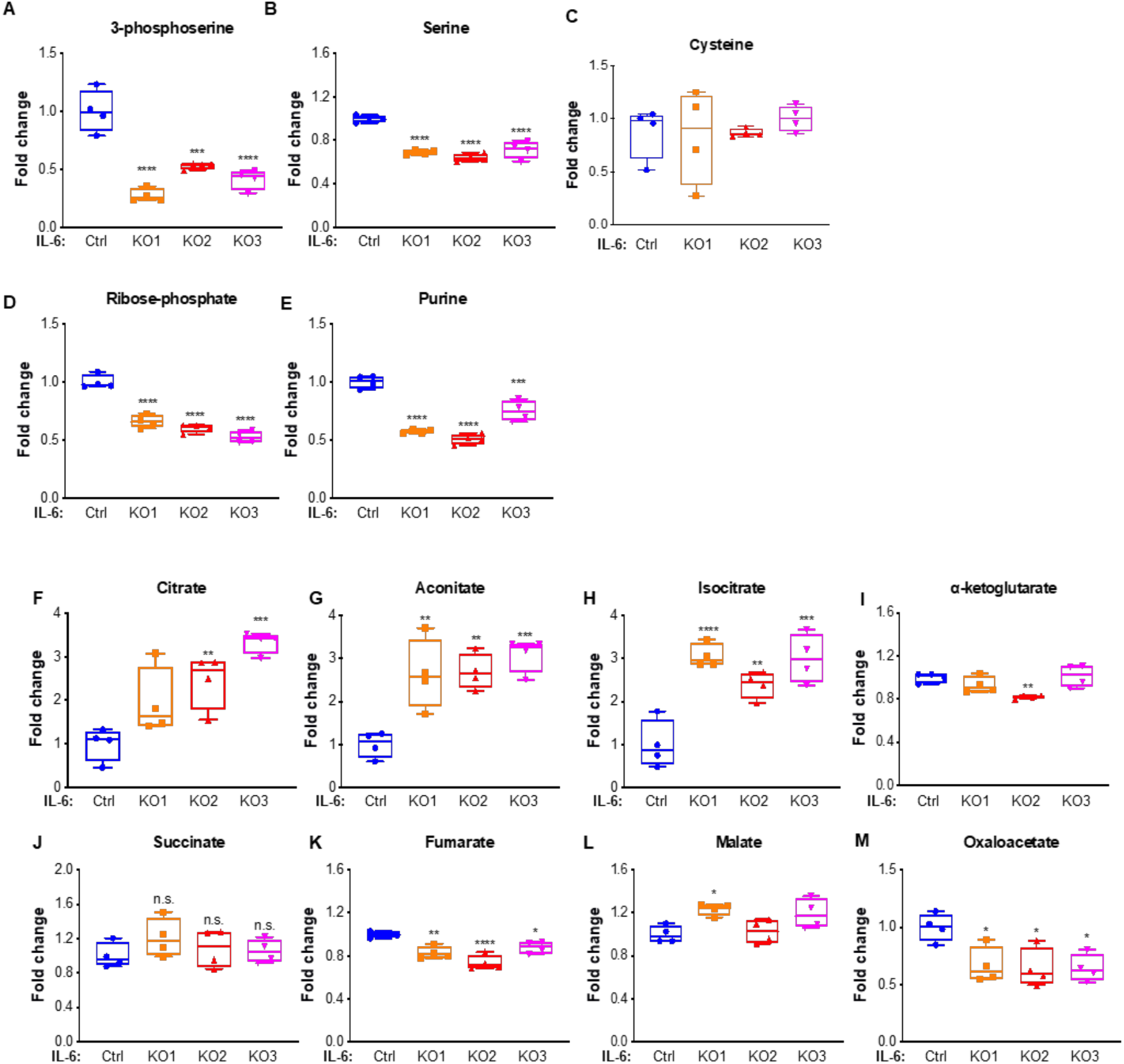
IL-6 knockout decreases serine in TSC2-deficient cells. (*A-M*) Metabolites of *de novo* serine biosynthesis, pentose phosphate pathway and TCA cycle quantified by LC/MS were differentially regulated in IL-6 knockout, TSC2-deficient MEFs compared to controls. Data presented as individual values with box and whisker plot showing mean ± minimum and maximum value of four biological replicates. Statistical analysis was performed by One-Way ANOVA *p<0.05, **p<0.01, ***p<0.001, ****p<0.0001 as compared to control.

**Fig. S5.**
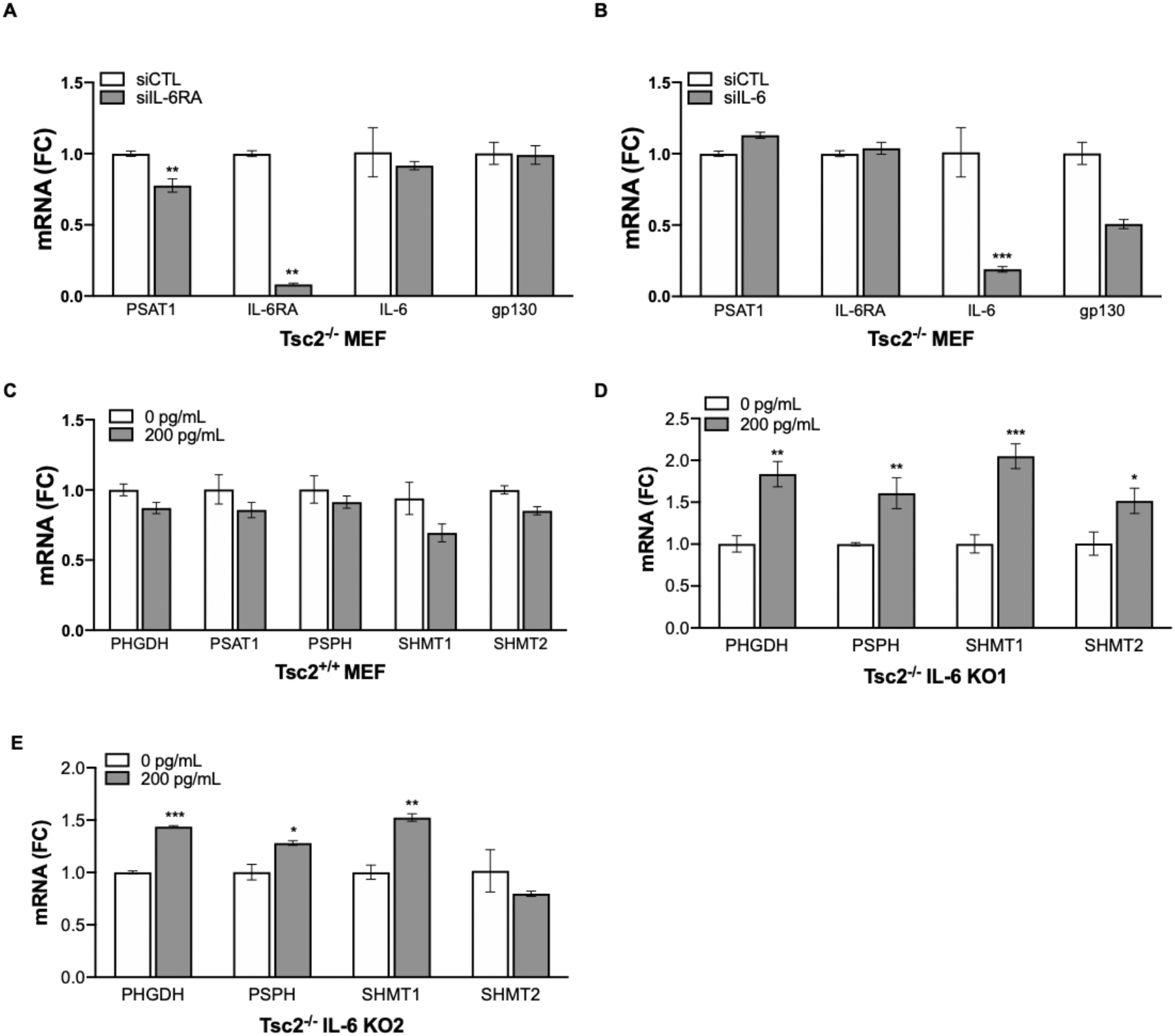
IL-6 modulates serine biosynthesis genes in a TSC2-dependent manner. (*A*) IL-6 receptor knockdown using siRNA decreased *PSAT1* mRNA expression in TSC2-deficient cells. (*B*) *PSAT1* mRNA expression is unchanged by IL-6 knockdown using siRNA in TSC2-deficient cells. (*C*) Recombinant IL-6 treatment has no impact on *de novo* serine biosynthesis genes in Tsc2 wildtype cells (IL-6; 200pg/ml; 24 hours). (*D* and *E*) Recombinant IL-6 treatment of IL-6 knockout, Tsc2-deficient cells increased the mRNA expression of serine biosynthesis genes (IL-6; 200pg/ml; 24 hours). The data are presented as mean ± SD of three independent experiments. Student’s t test was used for statistical analysis with * p<0.05, ** p<0.01, \*\*\**p*<0.001.

**Fig. S6.**
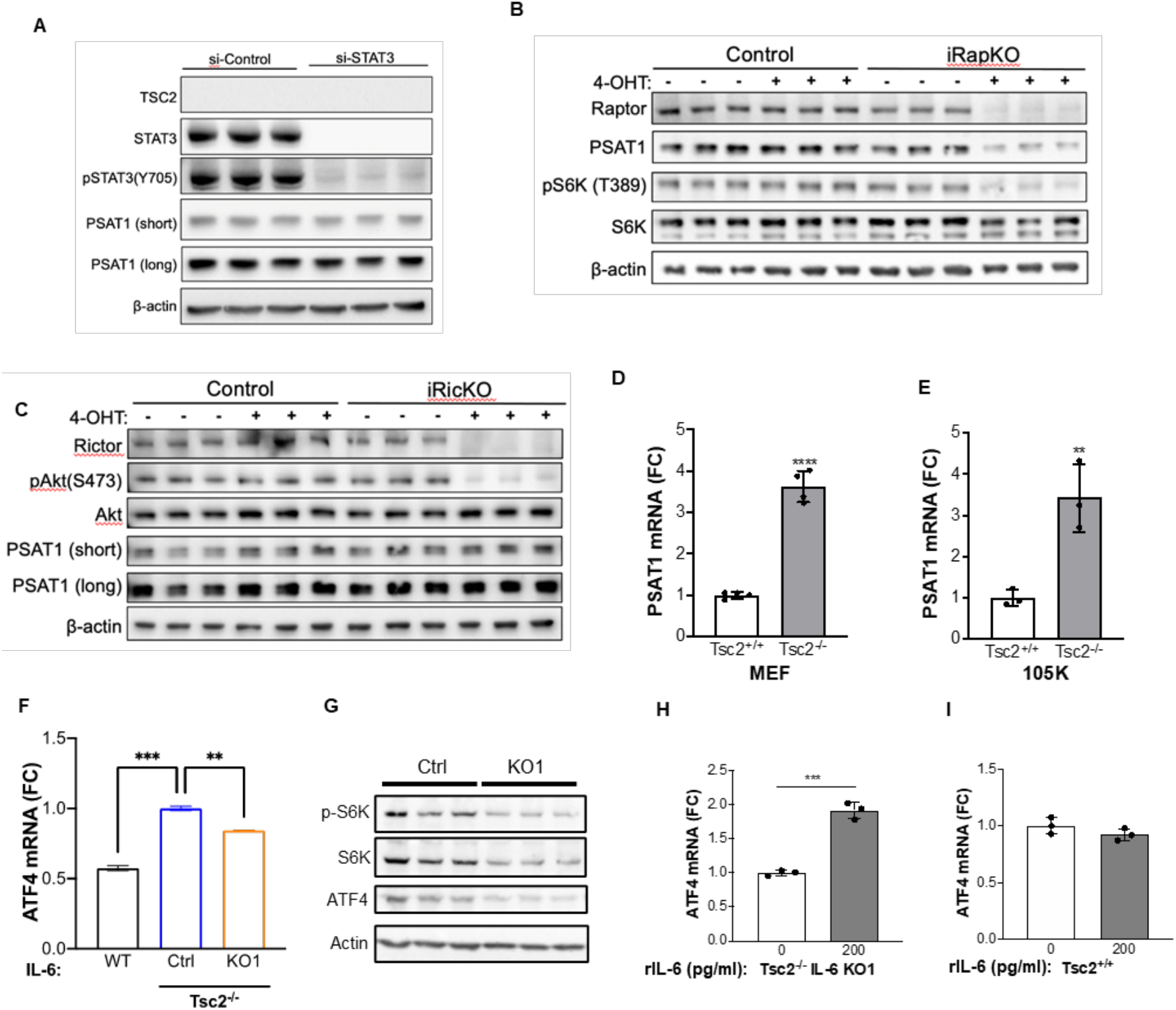
PSAT1 is regulated in a STAT3-independent mTORC1-dependent manner. (*A*) Western blotting showing that inhibition of STAT3 using siRNA did not impact PSAT1 protein expression (72h siRNA, serum-free last 24h). (*B*) mTORC1 inhibition using inducible Raptor knockout cells decreased PSAT1 protein levels. (*C*) mTORC2 inhibition using inducible Rictor knockout had no impact on PSAT1 protein levels. (*D*) *PSAT1* mRNA levels are increased in TSC2-deficient MEFs compared to TSC2-expressing MEFs. (*E*) *PSAT1* mRNA levels are increased in TSC2-deficient mouse kidney cystadenoma 105K cells compared to TSC2 re-expressing cells. (*F*) *ATF4* mRNA levels are increased in TSC2-deficient cells compared to TSC2-expressing cells, while IL-6 knockout decreased *ATF4* expression. (G) Knockout of IL-6 inhibited phosphorylation of the mTORC1 target gene S6 kinase and the protein levels of ATF4. (*H*) Recombinant IL-6 increased the mRNA expression of *ATF4* in IL-6 knockout TSC2-deficient cells (rIL-6; 200 pg/ml; 24 hours). (*I*) Recombinant IL-6 did not impact the expression of *ATF4* in TSC2-expressing cells rIL-6; 200pg/ml; 24 h). Data are presented as mean ± SD of three independent experiments. Student’s t test and One-Way ANOVA were used for statistical analysis with **p<0.01, ***p< 0.001, ****p<0.0001.

**Fig. S7.**
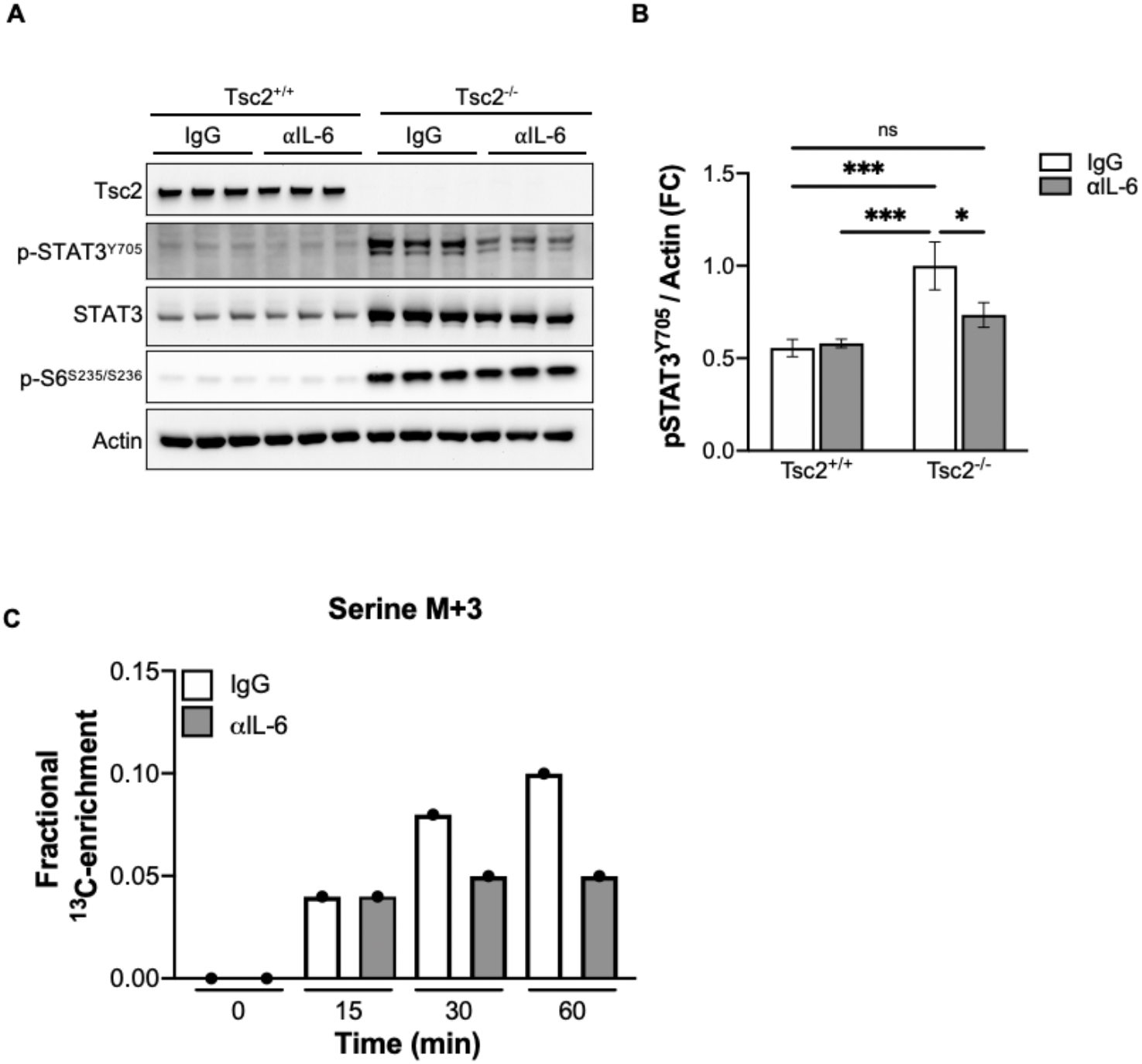
αIL-6 antibody reduced STAT3 signaling and serine synthesis in TSC2-deficient cells. (*A*) Western blot and (*B*) densitometry show that αIL-6 antibody decreased phosphorylation of STAT3^Y705^ in TSC2-deficient cells (αIL-6; 10ug/ml; 24 h). (*C*) U-^13^C glucose tracing following αIL-6 antibody decreased M+3 serine levels compared to IgG antibody control (10ug/ml). Data in C are derived from a single biological replicate for each treatment condition and time point.

**Fig. S8.**
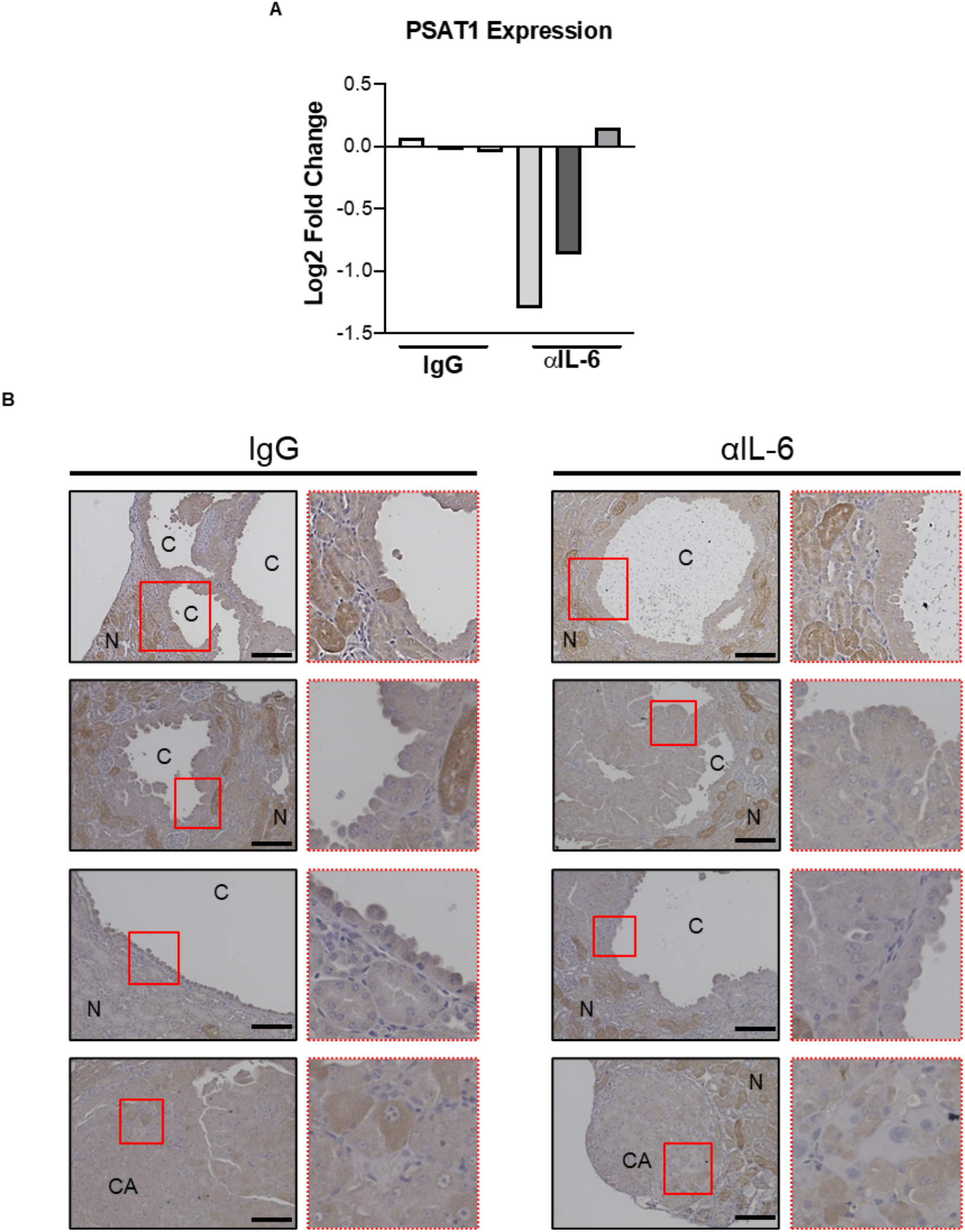
PSAT1 expression is reduced in renal lesions of Tsc2^+/-^ mice treated with αIL-6 antibody. (*A*) Semi-quantitative analysis of PSAT1 staining in three IgG and three αIL-6 antibody treated mice (200 ug/mouse, three times/week). (*B*) Representative PSAT1 staining from IgG treated kidney with outlined area in red enlarged in image (*Left column*). Representative PSAT1 staining from αIL-6 antibody kidney with outlined area in red enlarged in image (*Right column*). c-renal cyst, ca-cystadenoma, n-normal adjacent kidney, Scale bar = 100μm.

